# Multi-channel recordings reveal age-related differences in the sleep of juvenile and adult zebra finches

**DOI:** 10.1101/2022.06.08.495284

**Authors:** Hamed Yeganegi, Janie M. Ondracek

## Abstract

Despite their phylogenetic differences and distinct pallial structures, mammals and birds show similar electroencephalography (EEG) traces during sleep, consisting of distinct rapid eye movement (REM) sleep and slow wave sleep (SWS) stages. Studies in human and a limited number of other mammalian species shows that this organization of sleep into interleaving stages undergoes radical changes during lifetime. Do these age-dependent variations in sleep patterns also occur in the avian brain? Does vocal learning have an effect on sleep patterns in birds? To answer this question, we recorded multi-channel sleep EEG from juveniles and adult zebra finches for several nights. Whereas adults spent more time in SWS and REM sleep, juveniles spent more time in intermediate sleep (IS). The amount of IS sleep was significantly larger in male juveniles engaged in vocal learning compared to female juveniles, which suggest that IS sleep could be important for learning. In addition, we observed that the functional connectivity increased rapidly during maturation of young juveniles, and was stable or declined at older ages. Synchronous activity during sleep was larger for recording sites in the left hemisphere for both juveniles and adults, and generally intra-hemispheric synchrony was larger than inter-hemispheric synchrony during sleep. A graph theory analysis revealed that in adults, highly correlated EEG activity tends to be distributed across fewer networks that are spread across a wider area of the brain, whereas in juveniles, highly correlated EEG activity is distributed across more numerous, albeit smaller, networks in the brain. Overall, our results reveal significant changes in the neural signatures of sleep during maturation in an avian brain.

## Introduction

Sleep is a reversible state characterized by a reduced level of behavioral activity and responsiveness to the environment. Sleep - or a sleep-like state - has been observed across animal taxa, including early animals without a centralized nervous system, such as the cnidarian.^1, 2^

Reconstructing the evolution of sleep across groups of amniotes has been a major thrust of comparative sleep research, and these investigations have usually provided 1) a description of the presence or absence of behavioral sleep, 2) a search for electrophysiological evidence of SWS and REM sleep, and 3) a final interpretation about what the presence or absence of certain features of sleep in an animal means to the evolution of sleep within amniotes.

As major features of sleep were first defined in mammals, the search for electrophysiological features of sleep in both avian and non-avian reptiles was heavily influenced by mammalian-like indicators of sleep, i.e., the presence of slow waves or delta power, (indicative of SWS in the mammalian cortex), and non-SWS characterized, by low amplitude brain activity, concomitant eye movement, and decreased muscle tone (measured using electromyogramy (EMG) and indicative of mammalian REM sleep).

Pigeons were one of the first non-mammalian animals whose sleep was investigated,^3, 4^ and it is now generally accepted that birds undergo behavioral sleep and show signs of both SWS and REM sleep. In birds, SWS is characterized by delta activity (1-4 Hz) with some tonic muscle tone, and REM sleep is generally characterized by low amplitude brain activity and conjugate and occasional non-conjugate eye movements. Additionally, birds show an intermediate stage (IS) of sleep, which has been compared to the early stages of mammalian non-REM (NREM) sleep and which can be considered a transitional stage of sleep.^5, 6^

Despite these similarities, clear differences do exist between mammals and birds. For example, many birds remain vigilant during sleep, opening one or both eyes during SWS, and only closing both eyes during REM sleep.^7–9^ Sleep spindles, a hallmark of mammalian NREM, have not been observed during avian sleep.^5, 6, 10^ Furthermore, particular sleeping postures differentially affect nuchal muscle tone, creating difficulties in relying on clear differences in EMG as a definition of REM sleep versus SWS sleep states.^11, 12^ Finally, how should one interpret the behavior of migrating birds, which can fly continuously for days? Using a strict behavioral definition of sleep, flying birds cannot be sleeping (see ^13^ for an examination of how migrating birds sleep).

Although there are more than 10,000 species of birds,^14^ electrophysiological signs of sleep have been investigated in only a handful of these animals: chickens and pigeons;^3^ geese;^15^ white-crowned sparrow;^16^ zebra finch;^5^ burrowing owl;^7^ ostrich;^12^ sandpiper;^17^ and the tinamou,^18^ to name a few. The sleep studies of birds so far have focused primarily on adult birds, and the few studies of immature birds ^19–21^ do not provide a comprehensive understanding of how aspects of sleep change as a function of aging - or learning.

Vocal learning in songbirds is one of the few innate learning models that can be used to study the effects of sleep on memory in birds. As a juvenile bird learns to sing, it must undergo a complex memory task that involves the formation of auditory memories, sequences of motor output, and associative higher-order representations of learned vocalizations.^22^ The process of song learning is similar to speech learning in humans and consists of 1) auditory encoding of the song template during a sensory phase, and 2) the process of vocal imitation during the sensorimotor phase.^23, 24^ As soon as a juvenile bird is exposed to the song of another male bird (usually its father, the “tutor”), the young bird begins to imitate aspects of those songs in squeaky and noisy subsongs, which are often compared to the babbling of human babies.^25, 26^ Through a process of auditory feedback and motor learning, juvenile subsongs transition from acoustically simple songs to complex and stereotypical adult songs in a process known as crystallization.^27^

Considering the extensive alterations in sleep architecture that occur during development and across aging in mammals^28^ as well as the paucity of information available about sleep in juvenile birds - and specifically juvenile birds engaged in vocal learning - it is of major interest to characterize sleep across different ages of birds to examine whether there are clear differences between sleep features in adult and juvenile zebra finches.

## Results

### Sleep traces are globally similar across electrode sites

In order to investigate the differences between sleeping brain activity in juvenile and adults, we chronically implanted 10 zebra finches (3 adults; 7 juveniles; see Table 1 for more information) with custom-designed electrodes to record supra-dural EEG from 16 sites over a wide area of the skull (approx. 7 mm × 8 mm; Fig. 1A). We recorded brain activity during 12-hour periods of natural sleep for 49 total nights across animals (more than 600 hours of data for all animals). During the tethered sleep recordings, animals assumed typical avian sleep postures, such as tucking the head in the feathers (Supplementary Fig. S1A, B). The body movement, as assessed with video recordings throughout the night, was generally low (Fig. 1B, Supplementary Fig. S1C).

**Figure 1.**
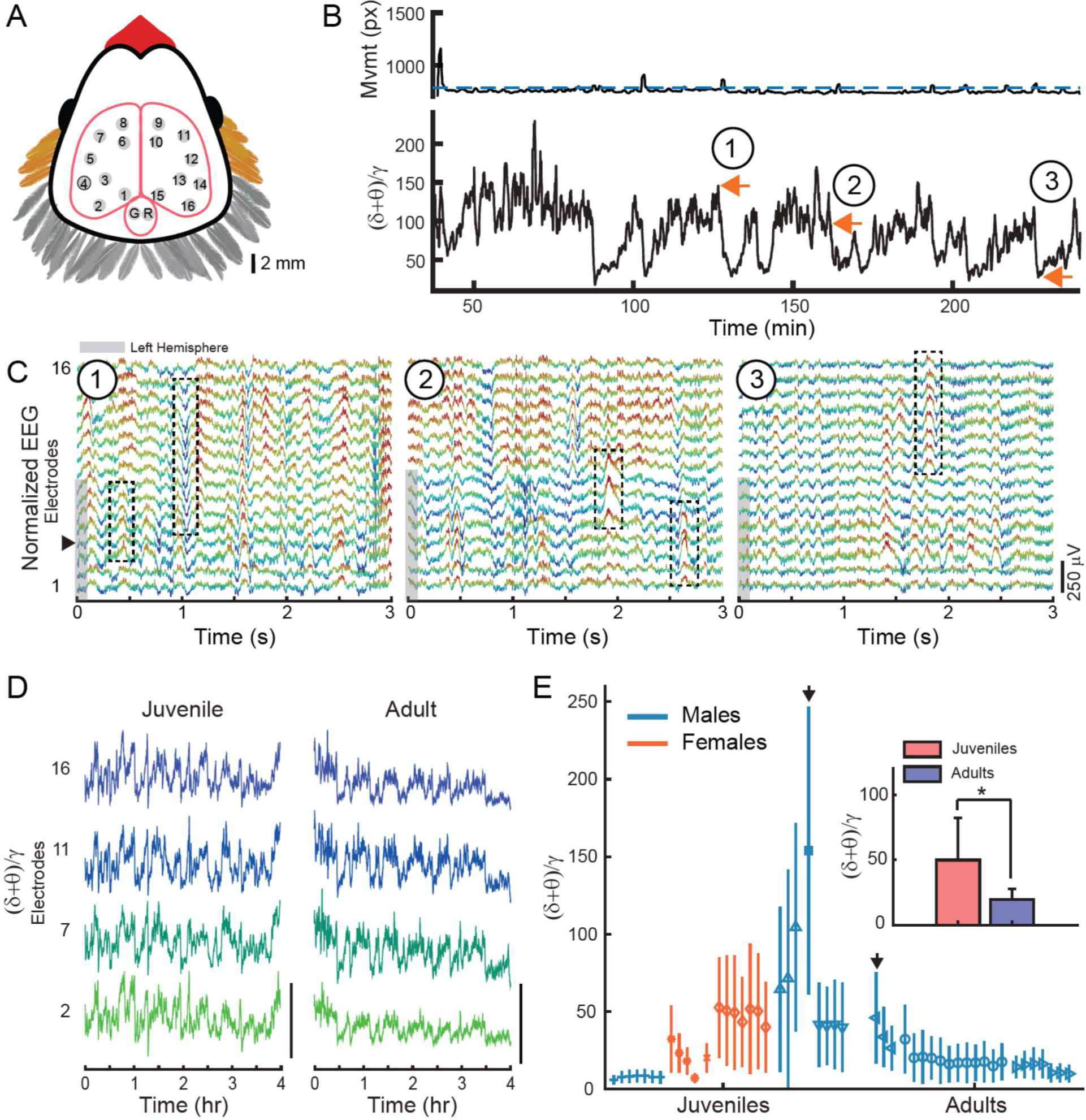
Multichannel EEG recordings during sleep. (A) Position of 16 EEG electrode sites on the surface of the skull: 8 electrode sites were located on each hemisphere and covered a wide area over the surface of pallium. Specific brain structures underlying each electrode are listed in Table 2. G, ground electrode; R, reference electrode. (B) Top: Bird movement extracted from the infrared video recording (px, pixels). Blue dashed line indicates the threshold delineating wake and sleep. Bottom: Corresponding (d+q)/g trace shows the oscillatory components of EEG, representing different sleep stages. Orange arrows 1, 2, and 3, correspond to EEG data in C. Movement and (d+q)/g traces are smoothed with a 30s window for visualization purposes. (C) 3s examples of simultaneous EEG recorded from the 16 different electrodes (bottom trace, electrode 1; top trace, electrode 16). Circled numbers correspond to orange arrows in B and indicate examples of SWS (1), IS (2), and REM sleep (3). Color scheme is for visualization purposes only. Gray shading on left indicates the electrodes that are located on the left hemisphere. Black dotted boxes highlight examples of local waves (see Fig. 3). Black arrow indicates electrode 4. (D) (δ+q)/g traces are similar across electrode sites in a time scale of minutes. Traces are for a single juvenile (left) and adult (right); exact nights are indicated with black arrows in (E). Scale bar is 200 (unitless (d+q)/g ratio). (E) Median (d+q)/g values calculated across 12 hours of sleep in 3s bins, for all nights of each bird (median ± interquartile). Each symbol represents a different bird, and different nights of sleep are presented sequentially. Teal lines indicate males; orange lines indicate females. (d+q)/g values are relatively consistent across nights within each individual bird, but large variability exists across different birds. Black arrows indicate animals and nights use in (D). Inset: (d+q)/g values averaged across all juveniles (red) and adults (blue). Error bars indicate the SD. A significant effect of age was present (p=0.032; two-way unbalanced ANOVA).

**Table 1.** Description of experimental birds used in this study. Bird name, symbol, sex, age in days post hatch (dph) at the first night of recording, and number of non-noisy electrodes recorded in the zebra finch is indicated. Table follows the same order as the data displayed in Fig. 1E.

| Name | symbol | sex | age (dph) | # of good electrodes |
| --- | --- | --- | --- | --- |
| w009 | + | m | 50 | 15 |
| w016 | * | f | 50 | 13 |
| w020 | □ | m | 53 | 16 |
| w018 | X | f | 51 | 14 |
| w021 | ◇ | f | 52 | 13 |
| w043 | △ | m | 69 | 15 |
| w041 | ▽ | m | 69 | 15 |
| 73-03 | < | m | 686 | 16 |
| 72-00 | ○ | m | 764 | 14 |
| 72-94 | > | m | 780 | 15 |

For each night, we compared the structure of sleep in adults and juveniles by determining the ratio of low frequency oscillations to gamma oscillations, calculated as the ratio of the relative power of the delta (δ) and theta (θ) bands (δ+θ = 1.5-8 Hz) to the gamma (γ) band (γ = 30-49 Hz).This metric is analogous to those used in mammals^29, 30^ and reptiles^31^ but accounts for the frequency bands prevalent in avian EEG.^5, 32^

We observed that across animals, the (δ+q)/g ratio generally varied on the order of tens of seconds (Fig. 1B, Supplementary Fig. S1C). We examined the raw EEGs at high, middle and low points of the (δ+q)/g ratio. High ratio values corresponded to EEG patterns typical of SWS, and low ratio values corresponded to REM sleep (Fig. 1C; Supplementary Fig. S1D). Middle (δ+θ)/γ values corresponded to an intermediate stage (IS) of sleep which has been compared to stage 1 and 2 of non-REM sleep in human.^6^

We examined whether the (δ+θ)/γ ratio changed as a function of anatomical position by comparing the (δ+θ)/γ traces for different quarters of the brain (Fig. 1D). We found that the (δ+θ)/γ ratio captured global aspects of sleep and was largely similar across a wide range of electrode sites (Fig. 1D). The correlation between (δ+θ)/γ traces computed across all electrodes for all birds was high (correlation coefficient = 0.64 ± 0.085; mean ± SD; see Supplementary Fig. S2A for individual correlations). Therefore, for the rest of the analysis in this work, we used electrode site 4 for all birds (Fig. 1A, black circle; Fig. 1C, black arrow).

### (δ+θ)/γ values are larger in juveniles compared to adults

We calculated the median (δ+θ)/γ value for each 12-hour night of sleep for each bird (Fig. 1E). Interestingly, the variability around the median was largely similar for different nights of sleep within the same animal, but this variability differed widely across animals. This finding mirrors results from studies of human sleep, where distinct inter-individual differences in EEG have been observed in human.^33^

When comparing median (δ+θ)/γ value of each night between adults and juveniles, we found that the (δ+θ)/γ values in juveniles were larger compared to the adults (Fig. 1E inset; (δ+θ)/γ = 49.9 ± 32.2 (mean ± SD) in juveniles versus 19.5 ± 8.4 in adults; n = 27 nights of sleep in juveniles and n = 22 nights of sleep in adults). To statistically assess the effect of age group, i.e. juvenile versus adult, as well as sex, we used an unbalanced analysis of variance (ANOVA). The ANOVA test revealed that the (δ+θ)/γ values were significantly different across age groups (p=0.032, two-way unbalanced ANOVA). The ANOVA did not show a significant effect of sex on the (δ+θ)/γ values (p=0.645, two-way unbalanced ANOVA); however, this is difficult to interpret as the adult group only consisted of male birds.

The larger (δ+θ)/γ values that we observed in juveniles could be due to an increase in the low δ+q frequencies or a decrease in the γ frequencies compared to adults. We examined the power in each of these bands separately (Supplementary Fig. S2B). Although the low δ+q frequencies were higher in juveniles compared to adults (Supplementary Fig. S2C; 0.088 ± 0.059 in juveniles versus 0.062 ± 0.080 in adults; mean ± SD), this difference was not significant (p=0.07, two-way unbalanced ANOVA). The γ power was not significantly different between juveniles and adults (Supplementary Fig. S2D; 0.0040 ± 0.0050 in juveniles versus 0.0038 ± 0.0052 in adults; p=0.47, two-way unbalanced ANOVA). The 2-way ANOVA did not indicate a significant effect of sex on low or high frequency power (p=0.18 in both cases).

These findings likely result from the fact that there was large variability in the normalized δ+q or γ frequencies across nights for some animals (see Supplementary Fig. S2B, adult bird symbol ‘right triangle’) which nevertheless resulted in stable and low (δ+θ)/γ ratios (Supplementary Fig. S2E).

### Adults spend more time in SWS and REM; male juveniles spend more time in IS sleep

In order to quantify the amount of time spent in each sleep state, we used a clustering-based sleep scoring method^5, 6^ to categorize sleep into discrete bins of REM sleep, SWS, or IS sleep. This allowed us to calculate the proportion and duration of each sleep stage, including the IS stage. We analyzed n=32 nights of sleep in this manner - the clustering-based algorithm did not converge to a plausible solution for 17 out of the 49 nights (Supplementary Fig. S3; Supplementary Fig. S4; see Methods for details).

At the beginning of the night, both adults and juveniles spent the largest amount of time in SWS (Fig. 2A). After about 6 hours of sleep, the amount of time spent in REM sleep gradually increased. This pattern of more SWS at the beginning of sleep and more REM sleep at the end of sleep is similar to the patterns reported in other animals^34, 35^ including adult zebra finches^5^ and budgerigars.^6^

**Figure 2.**
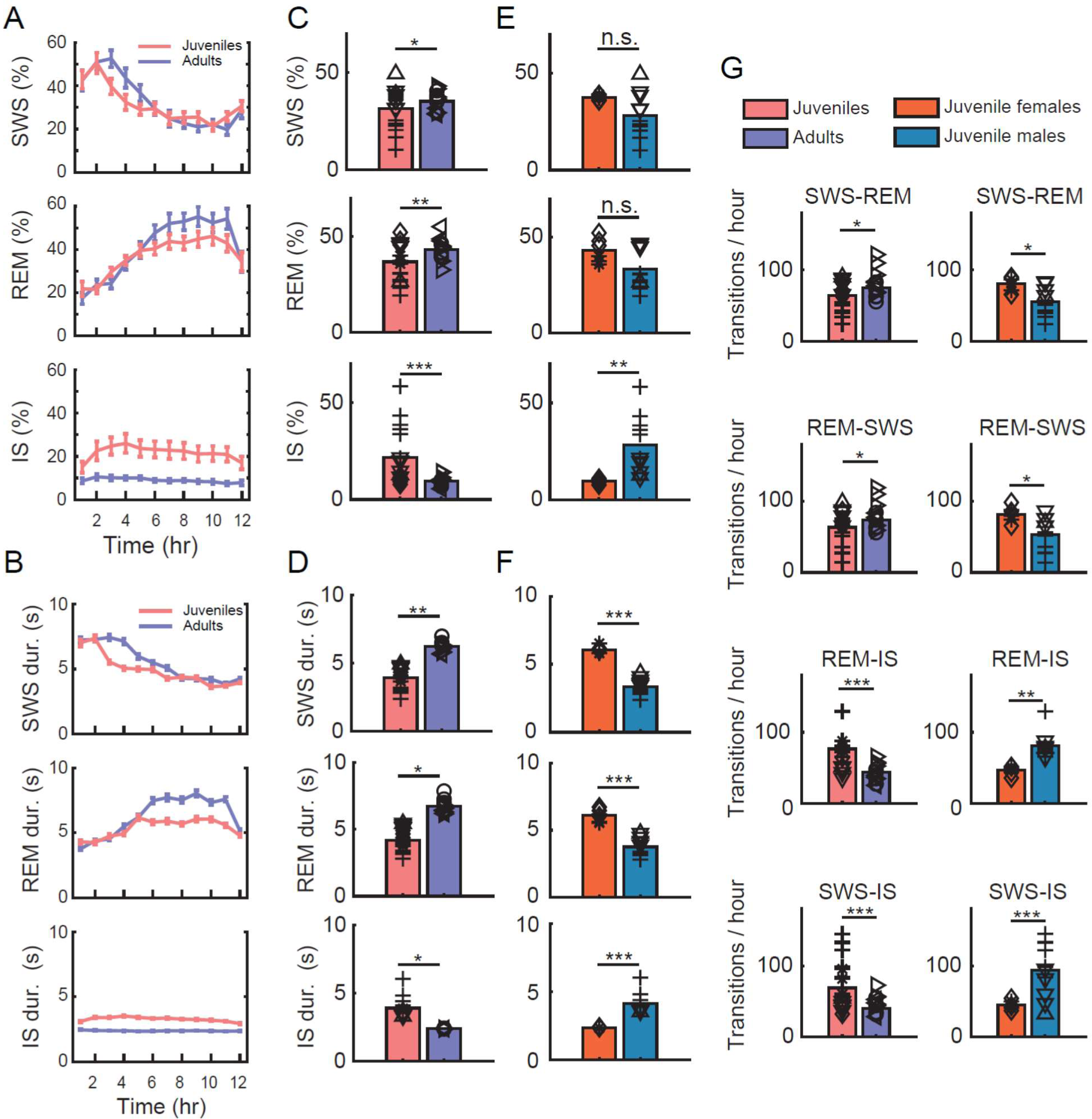
Sleep stage percentages and transitions in juveniles and adults based on automatic sleep segmentation. (A) Average percentage of SWS (top), REM (middle), and IS (bottom) for juveniles (red) and adults (blue) over 12 hours of sleep. Error bars indicate the s.e.m. Note how the early hours of the night are dominated by SWS, whereas the later hours of the night are dominated by REM for both adults and juveniles. (B) Duration of SWS, REM, and IS in juveniles and adults over 12 hours of sleep. Figure conventions same is in (A). (C) Mean percentages of SWS (top), REM (middle), and IS (bottom) pooled over all nights for juveniles (red) and adults (blue). Symbols indicate nights from individual animals (see Fig. 1E). Asterisks indicate significance (two-way ANOVA; * p ≤ 0.05; ** p ≤ 0.01; *** p ≤ 0.001). (D) Mean durations for SWS (top), REM (middle), and IS sleep (bottom) pooled over all nights for juveniles (red) and adults (blue). Figure conventions same as for (C). (E) Percentages of SWS (top), REM (middle), and IS (bottom) pooled over all nights for female juveniles (orange) and male juveniles (teal). Figure conventions same as for (C). (F) Mean durations for SWS (top), REM (middle), and IS sleep (bottom) pooled over all nights for female juveniles (orange) and male juveniles (teal). Figure conventions same as for (C). (G) Mean number of transitions per hour for juveniles (red) and adults (blue) and juvenile females (orange) and juvenile males (teal). Figure conventions same as for (C).

Similarly, for both adults and juveniles, SWS durations tend to be longer in the early hours of the night, whereas the REM sleep durations tend to be longer in the later part of the night (Fig. 2B).

On average, adult birds spent a significantly higher percentage of the night in SWS and REM sleep compared to juveniles (p=0.026 for SWS; p=2.7×10^-3^ for REM; two-way ANOVA; Fig. 2C). In contrast, juvenile birds spent significantly more time in IS sleep compared to adults (p=9.57×10^-6^; Fig. 2C). SWS and REM sleep durations were also significantly longer in adults, whereas IS durations were significantly longer in juveniles (Fig. 2D; SWS duration p=0.0018; REM sleep duration p=0.017; IS sleep duration p=0.014; two-way ANOVA).

We also observed a significant effect of sex on the SWS, REM, and IS sleep percentages (see Supplementary ANOVA statistics for p values), so we analyzed the juvenile males and juvenile females separately. While there was no significant difference for the percentage of the night spent in SWS and REM sleep between juvenile males and juvenile females, we found that males spent a significantly higher percent of the night in IS sleep compared to females (Fig. 2E).

We also observed a significant effect of sex on the SWS, REM, and IS sleep durations (see Supplementary ANOVA statistics for p values). Interestingly, female juveniles had significantly longer bouts of SWS and REM sleep, whereas male juveniles had significantly longer bouts of IS sleep (Fig. 2F).

We did not observe a negative correlation between the amount of time spent in REM sleep and age for juvenile zebra finches (Supplementary Fig. S5A), as reported elsewhere for owlets.^21^

### Juveniles transition to IS stages more frequently than adults

In addition to the amount of time spent in each sleep stage, we were curious as to whether there were differences in the transitioning patterns between sleep stages for juveniles and adults. Using the automatic sleep scoring method, we quantified all the SWS to REM sleep / REM sleep to SWS transitions as well as the SWS to IS sleep / REM sleep to IS sleep transitions (Fig 2G; see Supplementary Fig. S5B, C for transitions per hour). In general, adults transitioned significantly more frequently between SWS and REM sleep compared to juveniles (p=0.035 for both SWS to REM sleep and REM sleep to SWS transitions; two-way ANOVA), whereas juveniles transitioned significantly more to IS sleep stages (Fig. 2G; p=1.3×10^-7^ for SWS to IS sleep transitions; p=1.3×10^-5^ for REM sleep to IS sleep transitions; two-way ANOVA).

Similar to what we found for other sleep statistics, we also observed a significant effect of sex on the sleep transitions (see Supplementary ANOVA statistics for p values). We found that on average, female juveniles had significantly more transitions between SWS and REM sleep, whereas male juveniles had significantly more transitions to the IS sleep stages (Fig. 2G).

We also examined transitions using the continuous (δ+θ)/γ traces (Supplementary Fig. S6A). We focused on the transitions from SWS (high (δ+θ)/γ values) to REM sleep (low δ+θ)/γ values; Supplementary Fig. S6B; see Methods for details).

For adult birds, the transition frequency from SWS to REM sleep was relatively constant at around 45 transitions per hour throughout the night (Supplementary Fig. S6C; blue bars). In juveniles, however, the transition frequency varied throughout the night: it ranged from approximately 45 transitions per hour at the beginning of the night to about 30 transitions per hour in the middle of night, and rose to over 50 transitions per hour in the last hours of sleep (Supplementary Fig. S6C; red bars). For the 4th, 5th, and 6th hours of sleep, juveniles transitioned significantly less frequently between SWS and REM sleep compared to adults (Supplementary Fig. S6C; Wilcoxon rank-sum test, p=0.045, 0.001, and 0.0005 respectively).

We found that in juveniles, the variable frequency of transitions during the night was linked to the variable duration of time spent in SWS and the IS sleep phases (Supplementary Fig. S6B). When we examined SWS-REM-SWS transitions using high (δ+θ)/γ values versus low (δ+θ)/γ values, the average SWS cycle was significantly longer in juveniles compared to adults (Supplementary Fig. S6D; 92.86 s ± 121.41 s versus 82.84 s ± 102.80 s, unpaired t-test, p=5.7×10^-12^). Similarly, when we examined REM sleep cycles, the average inter-REM sleep interval was also significantly longer for juveniles compared to adults (92.88 s ± 115.07 s versus 82.74 s ± 99.15 s, unpaired t-test, p=5.5×10^-13^; Supplementary Fig. S6D).

Altogether, these results suggest that juveniles - and specifically male juveniles - may spend a longer period of time in IS sleep and transition to IS sleep compared to adults and female juveniles.

### Local waves occur more frequently during sleep in adults compared to juveniles

Sleep stages, such as REM sleep and SWS, are widespread events that are usually distinguishable based on their unique neural signatures.^3^ In both adult and juvenile zebra finches, we observed that global aspects of sleep were highly similar across recording sites (e.g., Fig. 1D). However, we also observed short duration (10-100 ms) “local waves” that spanned a narrow range of EEG electrodes (see Fig. 1C, black dotted boxes, for examples). We defined local waves as oscillations that occurred on 25% to 75% of the electrodes at a given time (see Methods for details). Importantly, these local waves were distinct from unihemispheric SWS activity^39, 40^ in which patterns of slow waves are observed globally in one hemisphere but not the other and which typically have durations of tens of second or longer.

We quantified the occurrence of local waves relative to the (δ+θ)/γ trace in both juveniles and adults (Fig. 3A, B). The occurrence of local waves was highly correlated with the (δ+θ)/γ values; that is, we observed that periods of high (δ+θ)/γ values corresponded to an increase in the number of local waves. The correlation between (δ+θ)/γ values and the occurrence of local waves, calculated across all nights in 3-s bins was significant and relatively high for the adults (0.52 ± 0.14; p=1.5×10^-19^; n = 22 nights; median ± interquartile interval, Pearson’s correlation test). The correlation between the (δ+θ)/γ values and the occurrence of local waves for juveniles, while lower, was also significant (0.30 ± 0.28; p=1.7×10^-4^; n = 27 nights; Pearson’s correlation test).

**Figure 3.**
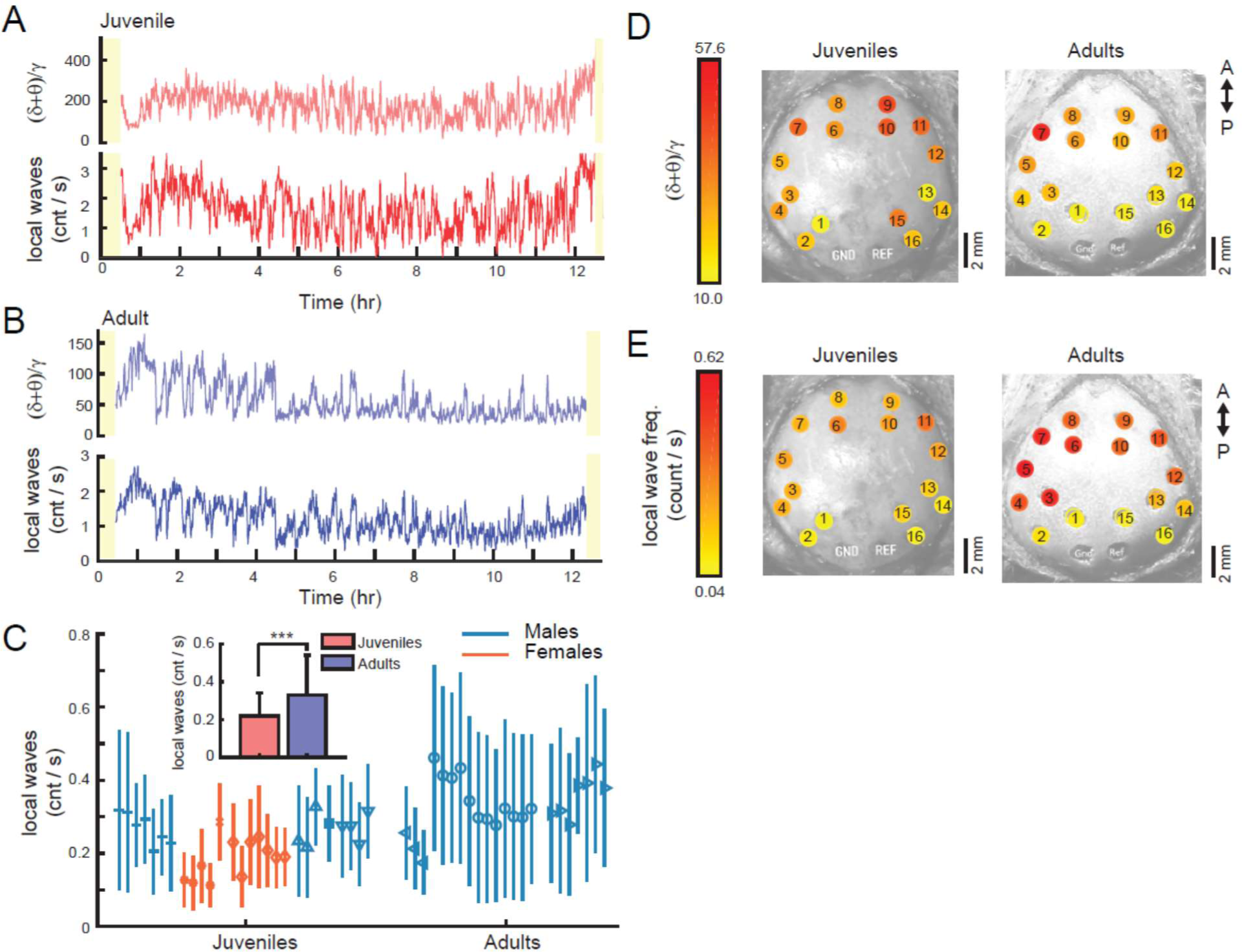
Local waves have a higher incidence in adults and a posterior-anterior axis. (A) An example of the (δ+θ)/γ trace for a juvenile bird (top, light red line) and the corresponding rate of occurrence for the local waves (bottom, dark red line) for a whole night of sleep (same juvenile as in Fig. 1D; black arrow in Fig. 1E). The occurrence of local waves was correlated with the (δ+θ)/γ ratio, i.e., local waves occurred more often during periods of high (δ+θ)/γ values. Note how this juvenile has a much larger median (δ+θ)/γ ratio compared to other juveniles and adults (see Fig. 1E), which may account for the large rate of local wave occurrence compared to the adult in (B). (B) An example of the (δ+θ)/γ trace for an adult bird (top, light blue line) and the corresponding rate of occurrence for the local waves (bottom, dark blue line) for a whole night of sleep (same adult as in Fig. 1D; black arrow in Fig. 1E). Data in (A) and (B) have been smoothed with a 30-s moving average filter for visualization purposes. (C) The average rate of occurrence for local waves across nights for all birds. Each symbol represents a different bird, and different nights of sleep are presented sequentially. Teal lines indicate males; orange lines indicate females. Vertical line indicates the SD for the night. Inset: local waves occur significantly more frequently in the adult birds (blue bars) compared to juveniles (red bars; p=6.7×10-7; two-way unbalanced ANOVA) Error bars indicate SD. (D) Average (δ+θ)/γ values computed at each electrode site across all juveniles (left) and all adults (right). (δ+θ)/γ values are higher in anterior regions (red colors) compared to posterior sites (yellow colors). This anterior-posterior gradient is especially pronounced in adults. GND, ground electrode; REF, reference electrode; A, anterior; P, posterior. (E) Average rate of local waves computed at each electrode site across all juveniles (left) and all adults (right). Same figure conventions as for (D). An anterior-posterior gradient is also present for the local waves, such that anterior sites have a higher rate of local wave occurrence compared to posterior sites.

In juvenile birds, the rate of local wave occurrence ranged from 0.19 to 0.32 events/s (Fig. 3C). When averaged over all the nights for all juvenile birds, local waves occurred at a rate of 0.22 ± 0.12 events/s (mean ± SD), or about 1 local wave every 4.5 seconds. The rate of occurrence was slightly higher in adult birds, and ranged from 0.28 to 0.46 events/s (Fig. 3C). When averaged over all adult birds, local waves occurred at a rate of 0.33 ± 0.21 events/s, or about 1 local wave every 3 seconds. This difference was significant (p=6.7×10^-7^; two-way unbalanced ANOVA; Fig. 3C inset). Furthermore, we also observed a small but significant effect of the bird’s sex on the local wave occurrence, such that local waves occurred significantly less frequently in females versus males (all males = 0.30 ± 0.19 events/s versus all females = 0.22 ± 0.14 events/s; p=0.013; mean ± SD; two-way unbalanced ANOVA).

### Local waves and slow waves are organized along an anterior-posterior axis during sleep

It is known that there is an anterior-posterior axis for slow wave activity in humans, such that larger slow waves are typically recorded on anterior EEG sites compared to posterior sites.^36^ We were curious to see if slow waves and local waves were similarly localized in the avian brain. Indeed, we observed that anterior EEG sites in both adult and juvenile zebra finches exhibited larger (δ+θ)/γ values compared to posterior positions (Fig. 3D; red colors). We observed a similar trend for the local waves, which occurred more frequently in anterior sites compared to posterior sites, especially for adults (Fig. 3E; red colors).

The anterior electrode sites were located above the hyperpallium (electrodes 8, 9, 6, 10; Fig 3D, E) and the lateral end of the mesopallium (electrodes 7 and 11), whereas the posterior sites were located above the song nucleus HVC (electrodes 1 and 15 in males), the end of the arcopallium (electrodes2, and 16), or mesopallium (electrodes 3, 13). These findings suggest that there could be a developmental aspect to the localization of local waves to anterior sites, such that during the process of brain maturation and cortical specialization, the neuroanatomical development of the avian brain facilitates the spread of low-frequency oscillations (1.5-8 Hz) in anterior sites.

### Functional connectivity is highest during SWS and lowest during REM in adults

Functional connectivity refers to the statistical (inter)dependence of neural activity in spatially distinct brain areas.^37^ In particular, inter-areal synchronization of oscillations is thought to be important for normal brain function^38^ and may facilitate the flow of neural information between different brain areas.^39^ We explored the inter- and intra-hemispheric functional connectivity of juvenile and adult zebra finches using the between-electrode Pearson’s correlation coefficient^38^ (CC; Methods).

For adult birds, but not the juveniles, we found that the functional connectivity was significantly different as a function of sleep stage (p=0.003 in adults; and p=0.66 in juveniles; Friedman test). In adults, the functional connectivity was highest during SWS (CC = 0.52 ± 0.29; mean ± SD) and lowest during REM sleep (CC=0.49 ± 0.27) when averaged over all pairs of electrodes (n=2299 pairs).

### Left intra-hemispheric connectivity is higher during sleep in adults

During sleep in both adults and juveniles, we often observed strong synchrony of the EEG signal within a hemisphere, but also across hemispheres (see Fig. 1C; Supplementary Fig. S1D). We quantified this by comparing between-electrode CCs for all pairs of intra-hemispheric (L-L pairs and R-R pairs) and inter-hemispheric electrodes (L-R pairs) separately during the whole night of sleep.

For both adult and juvenile birds, the intra-hemispheric CCs were larger than the inter-hemispheric correlations. For adults, the average intra-hemispheric CC was 0.62 ± 0.16 for L-L pairs (mean ± SD) and 0.57 ± 0.14 for R-R pairs, whereas the average L-R inter-hemispheric CC was much smaller: 0.27 ± 0.15 (Fig. 4A, blue bars). This difference was significant across comparisons (p=1.44e-53; two-way ANOVA) and post-hoc tests revealed significant differences for all three comparisons (Fig. 4A, asterisks; see Supplementary ANOVA table for exact p values). Notably, in the adult group, the functional connectivity was significantly higher for electrode pairs in the left hemisphere compared to the right hemisphere (p=0.040, post-ANOVA Tukey-Kramer multiple comparison test).

**Figure 4.**
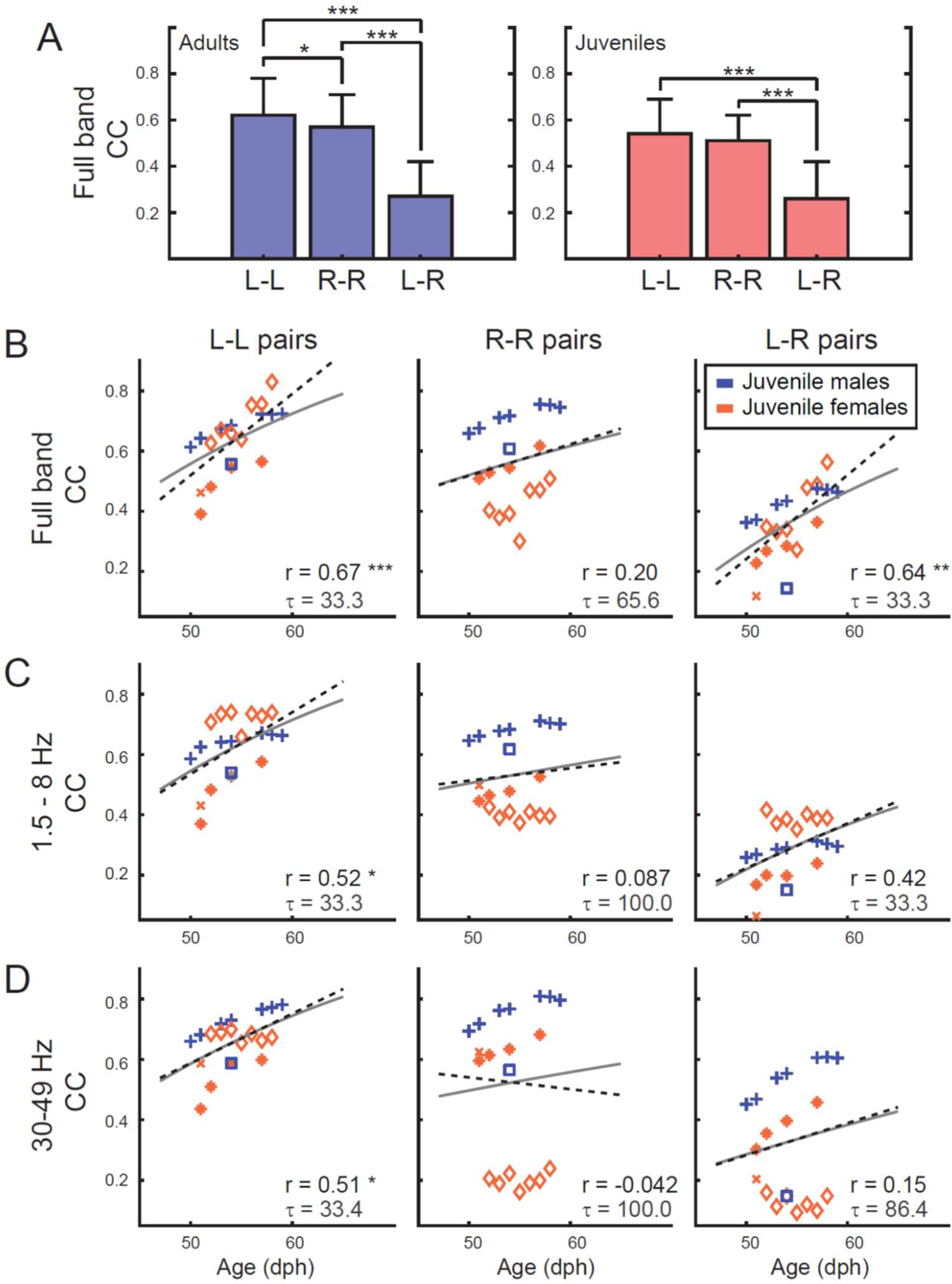
Left intra-hemispheric functional connectivity increases linearly during early maturation in juveniles. (A) Functional connectivity was calculated between pairs of electrodes using Pearson’s correlation coefficient (CC) on the full band EEG signal. Pair-wise comparisons of electrodes located in the left hemisphere (L-L) were calculated separately from electrodes pairs located in the right hemisphere (R-R). In both adults and juveniles, L-L and R-R CCs were significantly larger than CCs calculated for inter-hemispheric electrode pairs (L-R). Furthermore, in adults, L-L CCs were significantly larger than R-R CCs. (B) Scatter plot depicts the average L-L, R-R, or L-R CCs calculated for each night as a function of the juvenile birds’ age. Each symbol represents a different juvenile bird (same symbols as for Fig. 1E and Fig. 3C), and different nights of sleep are presented sequentially. Blue symbols indicate males; orange symbols indicate females. CCs were computed on the full band EEG signal (1.5 - 200 Hz). Black dotted line indicates the linear fit of the data, and the corresponding r value is located in the lower right corner. Asterisks indicate significant linear fits (* p ≤ 0.05; ** p ≤ 0.01; *** p ≤ 0.001). Gray line indicates the exponential fit of the regressed data, and the corresponding time constant τ is located in the lower right corner. A significant linear trend existed for L-L and L-R electrode pairs, but not R-R pairs. (C) Scatter plot depicts the average L-L, R-R, or L-R CCs calculated on the low frequency EEG band (1.5 - 8 Hz). Figure conventions same is in (B). For the low frequency band, only the L-L electrode pairs showed a significant linear increase in functional connectivity as a function of age. (D) Scatter plot depicts the average L-L, R-R, or L-R CCs calculated on the high frequency EEG band (30 - 49 Hz). Figure conventions same is in (B). For the high frequency band, only the L-L electrode pairs showed a significant linear increase in functional connectivity as a function of age.

The same trend was also true for juveniles (L-L = 0.54 ± 0.15; R-R = 0.51 ± 0.11; L-R = 0.26 ± 0.16 (Fig. 4A, red bars) and these differences were significant across comparisons (p = 9.64e-23; two-way ANOVA). However post-hoc tests revealed that only the inter-hemispheric comparisons were significant for juveniles (Fig. 4A, asterisks).

### Left intra-hemispheric functional connectivity increases linearly during early maturation in juveniles

We observed that the average intra- and inter-hemispheric CCs were higher in sleeping adults compared to sleeping juveniles, and we wondered whether we could observe an increase in the functional connectivity as a function of maturation. For this analysis, we used the data from the younger juveniles in our study (age < 60 dph, n=5 out of 7 juveniles; see Supplementary Fig. S7A, B for the data from other birds). We did not observe a significant change in the functional connectivity as a function of age in adult birds (p>0.05; t-test).

We used two methods to look for an age-related increase in the functional connectivity in juveniles. First, we analyzed whether there was a linear relationship between the intra- and inter-hemispheric CCs and the age of the bird. Second, we regressed the intra- and inter-hemispheric CCs as a function of the birds’ age (see Methods for details). To account for possible differences in the synchrony of the EEG oscillations at different frequencies, we computed the functional connectivity CCs for three different passbands: the full band (1.5-200 Hz; Fig. 4B), the low frequency band (1.5-8 Hz; Fig. 4C), and the high frequency band (30-49 Hz; Fig. 4D).

Interestingly, we observed that the functional connectivity increased linearly as a function of age for L-L electrode pairs in all frequency bands (Fig. 4C-D; R value asterisks), but that this was not true for R-R electrode pairs. Inter-hemispheric connectivity only increased as a function of age when computed for the full band, but not for the low or high bands individually.

When we examined the time constant τ arising from the regression model, values ranged from 33 to 100 days, demonstrating a fast increase of functional connectivity during maturation. We observed the highest rate of increase for the left hemisphere (L-L) pairs and for the inter-hemispheric (L-R) correlations.

These findings support the hypothesis that the juvenile zebra finch brain undergoes rapid changes in connectivity that stabilize in adulthood, and which is in line with observations in human studies.^40^

### Graph theory analysis reveals larger networks of correlated EEG activity in adults compared to juveniles

Although we observed broad differences in functional connectivity within and across hemispheres during sleep, we wanted to investigate functional connectivity at a finer spatial resolution. Borrowing methods from graph theory, (Fig. 5A; see Methods for details), we identified dominant co-active networks in juvenile and adults during SWS, REM sleep, and IS sleep. For each night, we considered each recording electrode as a “graph node” and analyzed 3-sec bins of EEG according to its putative sleep stage as determined by its (δ+θ)/γ value. We connected a pair of electrodes with a link if their activity was significantly correlated. Then we extracted the sub-networks for which each electrode was highly correlated with all the other electrodes in the sub-network. We only analyzed the sub-networks that appeared at least in 10% of the bins throughout the night, for three separate nights (See Supplementary Fig. S8 for results using an appearance threshold of 5% and 15%; results are in line with those presented here). For simplicity we denote such a highly correlated sub-network as a “dominant network”. Examples of these dominant networks are displayed in Fig. 5B (adult) and Fig. 5C (juvenile).

**Figure 5.**
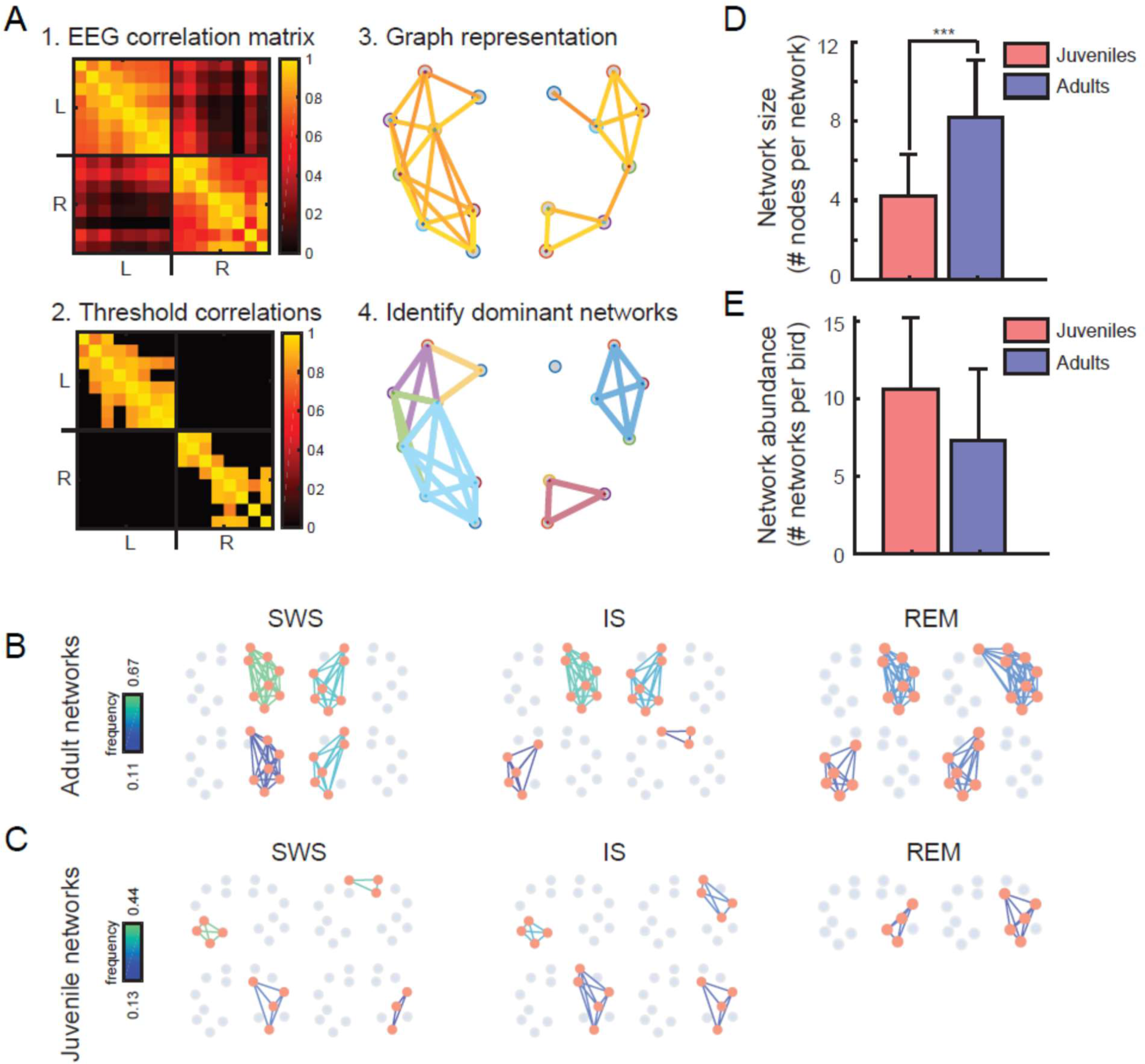
Highly correlated EEG activity is distributed across a few large networks in adults and several small networks in juveniles. (A) The main steps for the extraction of highly-correlated networks are illustrated (see Methods for more details). 1-The EEG correlation matrix was computed for each 3-sec bin. 2-Significantly correlated pairs were kept for further analysis, based on a statistical method46. 3-A graph representation of the significantly correlated pairs is constructed. To construct this graph, each EEG electrode was represented as a node, and a link connected two nodes in the case where the correlation between the corresponding EEG electrode was significantly high. 4. Using the Bron-Kerbosch algorithm from graph theory41 we extracted the sub-networks for which all electrodes were highly correlated, the “dominant networks”. (B) Dominant networks extracted from an adult during SWS, IS, and REM stages of sleep for 3 nights. A dominant network is indicated as a collection of orange nodes (the electrode sites) and the lines connecting them. Color coding of the lines represents the frequency of occurrence for each network across bins, i.e. the fraction of bins where the network appeared. Gray dots indicate nodes that are not included in the networks. In adults, dominant networks contain several nodes (are larger) but occur less frequently. (C) Dominant networks extracted from a juvenile during SWS, IS, and REM stages of sleep across all nights. Figure conventions same as in (B). In juveniles, dominant networks contain fewer nodes, but occur frequently. (D) Dominant network size was significantly larger for adults compared to juveniles. Error bars indicate the SD. (E) Dominant network abundance was lager for juveniles compared to adults, but not significantly different. Error bars indicate the SD.

We analyzed the 8 out of 10 birds for which we had at least 3 nights of data (3 adults and 5 juveniles). In these birds, the size of dominant networks (the number of nodes; Fig. 5D), and their abundance (the number of dominant networks per bird; Fig. 5E), was used as an indication of overall connectivity.

On average, the dominant network size was significantly larger for adults compared to juveniles (Fig. 5D; 8.2 ± 2.9 electrodes (mean ± SD) for the adults and 4.2 ± 2.1 electrodes for the juveniles; p=3.0 × 10^-8^, Wilcoxon rank-sum test). On the other hand, the networks were more numerous in juveniles compared to adults (Fig. 5E; 7.3 ± 4.6 networks on average for adults and 10.6 ± 4.6 dominant networks in the juveniles).

The graph theory analysis reveals two distinct patterns in adults and juveniles: in adults, highly correlated EEG activity tends to be distributed across fewer networks that are spread across a wider area of the brain, whereas in juveniles, highly correlated EEG activity is distributed across more numerous, albeit smaller, networks in the brain.

## Discussion

In this study, we analyzed the sleep EEG recorded in 12 hour periods from 3 adult and 7 juvenile zebra finches, totaling more than 600 hours of recorded data for all animals. We analyzed differences in the sleep patterns between juveniles and adults using two main approaches: 1) we analyzed sleep as a continuous variable using the (δ+θ)/γ ratio and 2) we categorized sleep into discrete stages using an automatic avian sleep clustering algorithm.^5, 6^ Whereas the former approach acknowledged the continuous nature of sleep-related dynamics, the latter approach allowed us to numerically characterize certain aspects of avian sleep and compare our results with other avian sleep studies.

We measured sleep EEG from 16 electrodes spanning the surface of the skull (Fig. 1A). We observed that across the electrode sites, (δ+θ)/γ traces were globally similar in the time-scale of minutes (Fig. 1D). Although we observed an anterior-posterior organization of (δ+θ)/γ values, such that anterior electrode sites tended to show larger (δ+θ)/γ values than the posterior sites (Fig. 3D), these observations suggest that it is sufficient to use 1 or 2 (anterior) electrodes to measure sleep EEG, as many avian sleep studies have done.^5, 6, 12, 13, 18, 21^

### Age-related differences in sleep between adults and juveniles: IS sleep and vocal learning

Vocal learning in songbirds is one of the few innate learning models that can be used to study sleep effects on memory. Although the link between REM sleep or SWS and vocal learning has not been investigated directly, behavioral experiments during song learning in juvenile zebra finches have highlighted a role for sleep.^42, 43^ Tutor song exposure causes a circadian pattern to develop in the vocalizations of the juvenile birds which continues into adulthood, but does not exist prior to tutor song exposure. Experiments in another songbird, the starling, using an operant conditioning paradigm to train birds to discriminate between different auditory stimuli, showed that sleep supports the consolidation of recent auditory memories.^44^

The juveniles in these experiments were housed with their fathers. During the time of the recordings, juvenile males were engaged in vocal learning, and juvenile females were exposed to their father’s songs and potentially engaged in auditory learning.^45, 46^

We observed several differences in the sleep patterns between adults and juveniles. When we segmented sleep into SWS, REM sleep, and IS sleep, we found that adults spent a significantly larger percentage of the night in SWS and REM sleep compared to juveniles, whereas juveniles spent significantly more time in IS sleep compared to adults (Fig. 2C). When we examined male and female juveniles, we found that the male juveniles spent a significantly larger percentage of the night in IS sleep compared to female juveniles and that IS sleep bouts were longer in duration for juvenile males compared to females. Furthermore, juvenile males transitioned significantly more frequently to IS sleep stages compared to females. These results suggest that IS sleep could be an important aspect of vocal learning in juvenile males.

IS sleep is distinct from REM sleep, and it often occurs as a transition state between SWS and REM sleep. IS sleep typically contains a mix of low-amplitude delta and higher-frequency elements such as theta^6^ (see Supplementary Fig. S3A for examples). As such, it is often not possible for human scorers to identify the onsets and offsets of IS sleep.^5, 6^ Indeed, analysis techniques that are reliant on human scorers to segment EEG data tend to categorize sleep into REM and NREM stages,^10, 47^ where NREM segments in this case contain both SWS and IS sleep, and the majority of the avian sleep literature focuses on REM versus NREM sleep.

Sleep in known to be important for learning, and specifically, SWS is known to have a critical role in the maintenance of memory in rodents^30, 48, 49^ and humans.^50–52^ However, it is unclear how to relate IS sleep to SWS. One commonality is that SWS and IS sleep are both are characterized by large, slow oscillations, and it may be that slow oscillations in general have a similar role in memory consolidation in the avian brain.^53^ Indeed, when we examined median (δ+θ)/γ ratios, we found that the (δ+θ)/γ ratios were significantly higher for juveniles compared to adults (Fig. 1E).

The role of slow waves and learning have not been thoroughly investigated in birds. One study in chicks observed a significant relationship between EEG theta oscillations that occurred during imprinting and memory consolidation during sleep, lending support to the idea that slow oscillations may have a role in avian learning.^54^ Additional investigations in birds that leverage behavioral paradigms such as vocal learning^55–57^ or spatial memory^58–60^ could help us better understand the interplay between slow oscillations and different types of memory in the developing avian brain.

### Sleep ontogeny in birds and mammals

Newborn mammals are thought to spend as much as 90% of their sleep time in a state that resembles an immature form of adult REM characterized by frequent muscle twitches, rapid eye movements, and irregular respiratory cycles.^61^ However, an alternative view suggested that REM sleep is actually not present at birth, but rather that these behavioral patterns could be spontaneous activity typical of an immature nervous system.^62^

In human development, children and adolescents spend more time in SWS. A meta-analysis of literature showed that the percentage of SWS is significantly negatively correlated with age.^63^ Similarly, a longitudinal study in children and adolescents revealed that NREM delta and theta activity drops sharply at the age of 10 and continues to decline until the age of 16.^64^ In humans, the reduction of slow waves is linked with reduction of gray matter during the course of cortical maturation in adolescence.^65^ We speculate that the importance of SWS during human adolescence might be similar to the importance of IS sleep for male juvenile zebra finches, as both phases involve intense phases of learning and neural reorganization. However, more studies that examine the exact role of IS sleep during learning in birds are required to explore this idea in greater detail.

Sleep ontogeny has not been thoroughly investigated in birds. One study examined sleep in young barn owls aged 27-48 dph and found that the amount of time spent in REM sleep decreased linearly with age.^21^ We did not find such a trend in our juvenile zebra finch data (Supplementary Fig. S5A). Rather, our work shows that adults have an increase in the amount of time spent in REM sleep and SWS and a decrease in IS sleep compared to juveniles. One obvious distinction between these two avian species is that juvenile zebra finches are engaged in vocal learning whereas juvenile barn owls are not. Similarly, zebra finches are daytime-active seed eaters, whereas owls are night-active predators. Such behavioral differences may account for the discrepancy in sleep patterns observed between these juvenile birds.

### The role of left hemisphere activity and brain lateralization in birds

Inter-areal synchronization of oscillations, also known as the functional connectivity, is thought to facilitate the flow of neural information between different brain areas. We found that in adult zebra finches, the functional connectivity was significantly higher in the left hemisphere compared to the right hemisphere (Fig. 4A). Furthermore, we found that in our young juveniles (n=5/7 birds) the functional connectivity increased linearly as a function of age for electrode pairs located on the left hemisphere (Fig. 4B). These results indicate an important role for neural synchronization of the left hemisphere. Indeed, several studies have already examined lateral asymmetry of the brain and behavior in the zebra finch and found left hemisphere dominance for both visual and auditory information processing related to courtship singing and visual displays.^66–68^

More generally, different cognitive tasks seem to be associated with different patterns of brain lateralization in birds.^69^ Larger left hemisphere activation has been observed in pigeons engaged in visual discrimination tasks, whereas larger right hemisphere activation has been associated with animals engaged in spatial memory tasks.^70^ In chickens, a task involving searching for food was associated with larger left hemisphere activation, while a predator avoidance task was associated with larger right hemisphere activation.^71^ These results suggest that there is an advantage inherent in the lateralization of certain tasks in birds. We observed that intra-hemisphere synchrony was higher than inter-hemisphere synchrony in both adults and juveniles (Fig. 4A). The synchronous (re)activation of brain areas during sleep might be an important aspect of the stabilization neural information during maturation and adulthood.

### An anatomical basis for reduced inter-hemispheric functional connectivity in birds

In this work, we found that the inter-hemispheric functional connectivity was almost half of the intra-hemispheric connectivity for both juveniles and adults (Fig. 4A). This result is almost certainly a function of the reduced anatomical pathways connecting the left and right hemispheres in birds.

In mammals, the corpus callosum heavily interconnects the right and left cortical hemispheres. The inter-hemispheric connections in birds are far fewer and the telencephalic pallium, sensory-motor, and associative regions receive limited input from the contra-lateral hemisphere.^72, 73^ The only inter-hemispheric pathways at the telencephalic level in birds consist of anterior commissure and the much smaller hippocampal commissure.^72^

In addition, the great majority of pallial areas do not participate by themselves in inter-hemispheric exchange in the avian brain.^72^ Instead, commissural exchange rests on a rather small arcopallial and amygdaloid cluster of neurons.^72^ In comparison, in the eutherian mammals which includes humans and rodents, the cingulate cortex and most of isocortex are connected to the contralateral homotopic regions via abundant interhemispheric corpus callosum tracts.^74^

However, it is worth noticing that the lower inter-hemispheric connectivity in the avian brain is not necessarily a disadvantage compared with placental mammals. For instance, unihemispheric sleep, which is observed in many bird species,^8, 9, 75–77^ including the zebra finch,^5^ helps the bird overcome the problem of sleeping in risky environment,^8^ and might be facilitated through weaker inter-hemispheric connectivity. Another indication of more independent hemispheres, that might be related to the absence of corpus callosum, is independent eye movements enabling birds to scan,^78^ and focus,^79^ upon different environmental stimuli simultaneously.

Since maintaining a normal level of functional connectivity is essential for the brain to function properly,^80–82^ we hypothesize that in birds, the evolution of brain connectivity took an alternative path in contrast with the emergence of corpus callosum in placental mammals; namely, that the left and right hemispheres developed stronger intra-hemispheric connectivity to compensate the lower inter-hemispheric connectivity while maintaining higher levels of hemispheric autonomy. However, more rigorous verification of this hypothesis would require further studies of functional connectivity in combination with brain stimulation to simultaneously measure effective connectivity, similar to human studies.^83, 84^ Further studies which examine functional connectivity in birds directly using non-invasive imaging techniques like functional MRI^85–87^ would be a welcome addition.

### The effects of courtship on sleep: Potential confounds and prospects for future research

In this study, juvenile and adult zebra finches were housed under different social conditions. We were not permitted to house animals in isolation, therefore, the 3 male adults were housed individually, but in visual and auditory contact with an adult female. In contrast, the male and female juvenile birds were housed individually, but in visual and auditory contact with their tutor father. Under these circumstances, the juvenile birds were exposed to the songs of their fathers and were assumed to be engaged in vocal learning (juvenile males) and/or auditory learning (juvenile females). In contrast, placing adult male birds in view of adult female birds might have induced courtship behavior.

This fundamental difference in the social housing conditions between adults and juveniles could have impacted the sleep in adults and juveniles differentially and could account for some of the differences we observed between juveniles and adults. For example, exposure to conspecifics of the other sex might modulate hormonal levels that affect arousal and activity during wakefulness, and which could in turn change aspects of sleep.

However, we observed that several aspects of adult male sleep matched the results from a previous sleep study in adult zebra finches,^5^ including 1) a decrease in SWS as the night progresses and 2) an increase in REM sleep as the night progresses. Furthermore, the durations of SWS, REM sleep, and IS sleep states for our adult birds were all within range of those values previously reported.^5^ This suggests that the differences we observed in sleep between adults and juveniles were not in fact due to differences in social housing. However further experiments, where for example the sleep EEG is recorded in adult females housed in visual and auditory contact with other adult females would help to clarify the potential effect of courtship behavior on sleep.

## Methods

### Ethical guidelines

All experiments were carried out in accordance with the principles of laboratory animal care and were conducted under the regulation of the current version of the German and European laws on Animal Experimentation (Approval # ROB-55.2-2532.Vet_02-18-108: J.M.O.), complying with ARRIVE guidelines. The study was approved by the ethics committee of the Government of Upper Bavaria (Regierung von Oberbayern; Committee following § 14.1 TierSchG (German animal welfare law)). Housing and breeding of animals were approved by the Veterinary Office of Freising, Germany (Approval # 32-568).

### Experimental animals

We used 3 adult and 7 juvenile zebra finches in these experiments. Zebra finches reach their sexual maturation at around 90 days post hatch (dph). The sex and the age of the birds at the first recording night are summarized in Table. 1.

Prior to experiments, birds were housed on site in aviaries and were kept on a 12:12-hr light: dark cycle. Finches received seed mix, millet, sepia bone, and water *ad libitum* and were given fresh salad and cooked egg supplements once a week.

### Behavior and video acquisition

Zebra finches were acclimated to the recording chamber for at least 5 days prior to the experiments. Experimental animals were kept together with a non-experimental female bird (adults) or the father tutor bird (juveniles) to prevent social isolation. Birds were separated from each other with a clear Plexiglas panel, such that animals could hear and see each other but could not interact physically.

In the recording chamber, finches received seed mix, millet, sepia bone, and water *ad libitum*. The recording chamber (total inner measurements: 120 cm × 50 cm × 50 cm) was equipped with LED lights, a UV bird light (Bird Systems, Germany), an infrared (IR) LED panel (850 nm), and had continuous air circulation. Video footage of the experimental animal was acquired with a near-IR sensitive camera (acA1300-60gm, Basler Ag, Germany) with custom-written software (ZR View, Robert Zollner) at a frame-rate of 20 fps, triggered with a pulse generator (Pulse Pal,^88^ Sanworks, NY, USA).

### Electrode preparation and positioning

For each experiment, we constructed silver ball EEG electrodes from silver wire (254 µm bare wire diameter, Science Products GmbH, Hofheim, Germany) by melting the tip of the wire into a smooth ball of approximately 0.50 mm in diameter using a Bunsen burner (CFH Löt- und Gasgeräte GmbH, Offenau, Germany). 18 electrodes were implanted: 8 electrodes were implanted over each hemisphere, and 2 additional ground and reference electrodes were implanted over the cerebellum (Fig. 1a). Electrode sites varied slightly across adult birds to avoid superficial blood vessels (variation less than 0.5 mm). The adult electrode map was rescaled for juveniles due to difference in head size, in order to maintain the relative electrode spacing. The brain structures underlying each electrode, according to the Nixdorf-Bergweiler zebra finch brain atlas^89^ are listed in Table 2.

**Table 2.** Position of the electrode relative to the underlying brain structures.

| Bilateral electrode pair | Underlying brain structures |
| --- | --- |
| 1, 15 | Medial end of HVc, Caudal end of Mesopallium |
| 2, 16 | Tractus dorso-arcopallialis, end of Arcopallium |
| 3, 13 | Lamina mesopallialis |
| 4, 14 | Lateral end of Lamina mesopallialis |
| 5, 12 | Hyperpallium apicale, lateral side of Lamina frontalis suprema |
| 6, 10 | Hyperpallium apicale, frontolateral side of Lamina frontalis suprema |
| 7, 11 | Lateral end of Mesopallium |
| 8, 9 | Frontal end of Hyperpallium apicale |

### Anesthesia and Surgery

Zebra finches were anesthetized with isoflurane (1-3 %) mixed with oxygen generated with an oxygen concentrator (EverFlo OPI, Phillips, Netherlands) and administered using a vaporizer (Isotec 4, Groppler, Germany) through a small tube directly into the beak of the bird. Excess isoflurane was collected and filtered (Scavenger LAS, Groppler, Germany). The body temperature was monitored with an infrared thermometer (EVENTEK E300, ShenZhen ShengYa Hardware Products Co, China), and maintained at above 39.5°C through the use of a homoeothermic heating pad (Harvard apparatus, MA, USA).

Under anesthesia, the experimental animal was positioned in a small animal stereotaxic frame (Kopf Instruments, CA, USA). The scalp was anesthetized with xylocaine (Xylocain Pumpspray, Aspen Pharma Trading Limited, Ireland), the feathers were removed with forceps, and the scalp was resected along the midline. For each EEG electrode, a small hole was drilled above the forebrain with a high speed dental drill (Volvere i7, NSK Europe GmbH, Germany). EEG electrodes were placed in the sites and secured using dental cement (Paladur, Henry Schein Dental, Germany).

Following surgery, the animal was released from the stereotaxic frame and administered a dose of Metamizol (100-150 mg/kg, i.m.). The animal remained on a heating pad until full recovery from anesthesia. An antibiotic (Baytril, 1025 mg/kg, i.m.) and an analgesic (Carprofen, 4 mg/kg, i.m.) were administered up to 3 days post-operatively. Following recovery from surgery, the animal returned to entirely normal feeding, sleeping, and singing behaviors as observed before surgery.

### Overnight electrophysiological recordings

Recordings were started at the second night after the implantation. One hour before lights off (light off period: 9:00 PM - 9:00 AM), the animal was connected to the head-stage and custom-made lightweight tether cable and allowed to assume a natural sleep posture on a branch that had been placed on the floor of the chamber. The camera was positioned to allow for the best visualization of the animal during sleep. The animal was allowed to sleep naturally overnight, and was disconnected from the headstage and tether cable shortly after lights turned on. Data from the first night after surgery was not used in the analysis.

The EEG electrodes were connected to a preamplifier (Intan Technologies, RHD2132), which was connected to the data acquisition board via the tether cable. The implant (cement, omnetics connectors, and wires) weighed ∼0.5-0.6 g. The preamplifier weighed 0.95 g. This combined weight of approximately 1.5 g was offset with an elastic string that was connected to the tether cable and was in the range of other impants and microdrives used in zebra finches.^90–92^ Recordings were performed with an Open Ephys acquisition board^93^ (Open Ephys Production Site, Portugal). Recordings were grounded and referenced against one of the reference wires. Signals were sampled at 30 kHz, wide band filtered (0.1-9987 Hz), and saved in a continuous format using the Open Ephys GUI.

### Anatomy

After the period of recording was over, the animal was deeply anesthetized with an overdose of sodium pentobarbitol (250 mg/kg, i.m.) until the corneal reflex disappeared. Afterward, the animal was decapitated, the brain was removed from the skull and post-fixed in 4% paraformaldehyde in PBS for postmortem verification of electrode sites.

## Data Analysis

### Video Motion Extraction

We developed a program in MATLAB (The MathWorks Inc., Natick, MA, USA) to extract a single variable that represents the overall amount of frame-by-frame movements. For each 2 consequent frames (*m* × *n* matrices containing gray-scale pixel values), we first computed the absolute value of the difference (equation (1)).

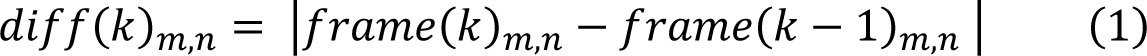

where *k* represents the frame number. *diff*(*k*)_*m,n*_ will have a non-zero value at any pixel that has changed from frame *k* − 1 to frame *k*. Then we extract the weighted average of *diff*(*k*)_*m,n*_ along the x and y axes separately. For example for the weighted average along the x axis, we sum up the *dif*(*k*)_*m,n*_ values along the rows and then scale it by the vector [1 2 … *n*], the x vector, (equation (2)).

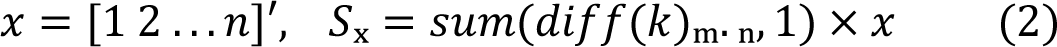

*S_y_* is acquired in the same way. At the end, using pythagoras formula, we merge Sx and Sy to achieve the total magnitude of movement (equation (3)).

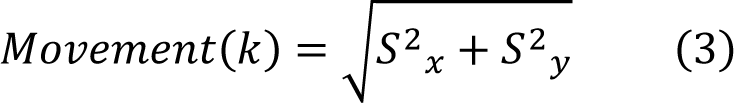

### EEG preprocessing and (**δ+θ)/γ** ratio analysis

EEG was down-sampled by a factor of 64 (final sampling rate: 468.75 sps). The EEG signal from each electrode was normalized (using the MATLAB “z-score” function) to avoid inhomogeneities that might arise from changes in the impedance of electrodes over days and binned into 3-s windows.

Sleep was behaviorally defined based on movement extracted from the video data. Bins where the movement was above its median by 2 interquartile intervals were labeled as awake and consequently excluded from analysis. This automatic sleep/wake labeling was validated on one data set that was that was also labeled visually as ground truth. Additional bins were excluded for which the EEG amplitude was more than 4 interquartile intervals above the zero base line. In total, approximately 400, 3-s bins (or 20 minutes of data, or less than 3%) were discarded per night of recording due to artifact rejection. Finally, some electrodes were broken and were therefore excluded from all further analysis. The number of used electrodes for each bird is reported in Table 1.

We first assessed the sleeping brain dynamics not in discrete stages, but as a continuous variable, the (δ+θ)/γ ratio, defined as the ratio of power in delta and theta bands (δ+q; 1.5 - 8 Hz) over the power in low gamma (γ; 30-49 Hz). Spectral power was computed using the Multitaper method with the time-halfbandwidth product equal to 1.25 and the number of DFT points equal to 2048.

### (**δ+θ)/γ** correlation for electrode pairs

In order to determine which electrode to use for the analysis, we computed the (δ+θ)/γ ratio in 3-s windows for each electrode separately for the first night of recording for each bird. The correlation coefficient between difference (δ+θ)/γ ratios was calculated between each electrode pair and averaged, resulting in a single value for each bird (Supplementary Fig. S2A. The average (δ+θ)/γ correlation for the different electrode pairs was quite high for each individual bird, and therefore we chose to use electrode 4 for our (δ+θ)/γ ratio analysis for each animal.

### Sleep stage identification based on the (**δ+θ)/γ** ratio

Previous research in zebra finch sleep shows that each stage of sleep, i.e. SWS, IS, or REM are computed based on their gamma- and delta-power content.^5^ That means, for example, SWS bins contain the highest delta power, REM the lowest, and the IS bins stand in between. Therefore, in order to include samples from different stages of sleep, we first ordered the EEG bins according to their (δ+θ)/γ values. The highest 10 percent were considered SWS, the lowest 10 were REM, and the 10 percent of the data encompassing the median (δ+θ)/γ values formed the IS group.

### Stage transition detection

In order to detect the number of transitions from SWS to REM, we used a moving threshold calculated over 30-minute windows of the non-smoothed (δ+θ)/γ values. The SWS threshold was defined as the moving average plus 0.4 × moving iqr (interquartile interval calculated over 30-minute windows), and the threshold for REM was defined as the moving average minus 0.4 × moving iqr.

### Clustering-based sleep staging

Apart from detection of SWS and REM, based on thresholding the L/G, as described above, we also adopted an automatic procedure to label each 3-second window of sleep as one of the IS, SWS, or REM stages. The algorithm we have used here is described previously^5, 6^ and has been used for segmenting sleep using budgerigar and zebra finch EEG. The main steps in this algorithms are as follows: 1. Sleep EEG data is binned into non-overlapping 3-sec windows, 2. Several stage-dependent EEG features, e.g. Delta power, are extracted, 3. Two clusterings for SWS vs non-SWS and REM vs non-REM are performed in parallel for all windows, and 4: The sleep stage label is decided based on the outputs of the two clustering algorithms from previous step. Finally, we visualized the sleep scoring labels along the EEG to verify if the automatically-assigned labels are conforming with the EEG patterns of each specific stage; in most of the cases (32 nights out of 49 recordings for n=5 juveniles and n=3 adults; see Supplementary Fig. S4 for more details) the sleep scoring with this method was valid and therefore used in this paper. On other nights the k-means clustering used in this scoring scheme converged to a numerical solution, which was a local minima of the cost function in k-means clustering, but did not yield acceptable scoring outputs (n=17 nights; see Supplementary Fig. S3 for examples of successful and unsuccessful clustering).

### Detection of local waves

Normalized EEG data was filtered using a band-pass filter (0.5 - 20 Hz) to suppress the high frequency components that might affect the precise localization of wave peaks and troughs. Local wave detection consists of two steps: (1) detection of all the wave peaks or troughs across all the electrodes, and (2) extraction of the simultaneously-occurring waves.

All positive or negative peaks that were more than one inter-quartile interval away from median were detected. To prevent the double-detection of waves with a negative deflection following the positive peak, or vice versa, we set a minimum inter-wave period of 90 ms where subsequent peaks were not detected. We only considered peaks detected across different electrodes which occurred within a 10 ms time widow as simultaneous. Finally, in order to be considered a local wave, a wave needed to be present in only a subset of all the electrodes. Here we considered the simultaneous observation of peaks across the electrodes as a local wave if the peaks were present in at least 4 electrodes (25%) and not more than 12 electrodes (75%). This requirement was not constrained to a particular hemisphere, and indeed, some local waves were detected across hemispheres (see Fig. 1C for examples).

### Connectivity analysis

In order to measure intra-hemispheric EEG correlation, we averaged the correlations between all pairs of right hemisphere electrodes (R-R; numbering 15, 21, or 28 pairs of electrodes, corresponding to 6, 7, or 8 non-noisy electrodes on the right hemisphere) and all pairs of left hemisphere electrodes (L-L; numbering 21 or 28 pairs of electrodes, corresponding to 7 or 8 non-noisy electrodes on the left hemisphere). The inter-hemispheric connectivity was computed as the average of the correlation between the pair of electrodes from opposing hemispheres (42-64 pairs, depending on the number of non-noisy electrodes). This correlation analysis was performed separately for the adult and juvenile group, and for 3 different frequency bands: low frequency (1.5-8 Hz), gamma (30-49 Hz), and full band (1.5-200 Hz).

### Regression Analysis

The juvenile CC values were fit with the following regression function:

Connectivity=A-B*exp(-age/τ),

where A and B are model parameters accounting for the initial point and the range of the exponential function and the time constant, τ, represents the time it takes for the connectivity to reach 63 % of the maximum, an index of growth rate.

### Graph theory analysis

We explored the compartmentalization of brain activity by extracting dominant networks of activity using methods from graph theory. We define a dominant network as a subset of EEG electrodes wherein (1) each pair of electrodes are significantly correlated and (2) it has the maximum number of such electrodes, i.e., no other electrode could be added that correlates highly with the current electrode s. For example, if there are 4 EEG electrodes which are all highly correlated with each other, and no other electrode exists which is highly correlated with all of these 4 electrodes, then these 4 electrodes form a dominant network. Such a network is called a “clique” in graph theory literature.

We extracted dominant networks for each stage of sleep separately. First, we computed the pair-wise correlation matrix for each 3-second bin belonging to the given sleep stage (e.g., REM, SWS, or IS) for all electrodes. From the correlation matrix, the statistically significant correlated pairs of electrodes were extracted using a statistical inference technique.^94^ A MATLAB implementation of this method was provided by the authors of ^94^. Having the significantly correlated pairs of EEG, the cliques were extracted using a MATLAB implementation of the Bron-Kerbosch algorithm.^41^ Among the obtained dominant networks, we discarded the ones involving fewer than 3 EEG electrodes and appearing in fewer than 10% of the bins for that sleep stage. In the end, for all the recording nights of a given bird, we kept the networks that were consistent across all the recording nights in a given sleep stage. The size of these dominant networks, i.e. number of EEG electrodes involved, and number of these networks were reported in Fig. 5D and Fig. 5E.

### Statistics

We have used two-way analysis of variance (ANOVA) to compare the (δ+θ)/γ values in juvenile and adults, while accounting for age and sex independently. Furthermore, we used a one-way ANOVA to compared sleep statistics between male and female juveniles. Sometimes, for the sake of brevity, some p values are reported only in the Supplementary Information ANOVA statistics. The complete table of statistics for each ANOVA test can be found in the supplementary information. In other cases, we use, t-tests, Wilcoxon ranksum test, or Friedman tests, where appropriate.

## Availability of Data and Materials

All data and analysis reported in this paper are available from the corresponding author upon request.

## Supporting information

Supplementary Information - ANOVA Statistics

## Acknowledgments

This research was funded by grants from the German Research Foundation (Deutsche Forschungsgemeinschaft; ON 151/1-1) and the Daimler and Benz Foundation (Postdoctoral scholarship) to J.M.O. The authors are grateful to B. Seibel and Y. Schwarz for technical assistance; C. Fink and E. Jochen for help with mechanical design, fabrication, and electronics; A. Schuhbauer for administrative assistance, and H. Luksch, U. Firzlaff, T. Fenzl, and M. Shein-Idelson, and two reviewers for feedback on previous versions of the manuscript.

## Author contributions

Conceptualization, J.M.O.; Methodology, J.M.O. and H.Y.; Software: H.Y. and J.M.O.; Validation, J.M.O.; Formal Analysis, H.Y.; Investigation, H.Y.; Writing – Original Draft, H.Y.; Writing – Review & Editing, J.M.O.; Funding Acquisition, J.M.O.; Resources, J.M.O.; Visualization, H.Y. and J.M.O.; Supervision, J.M.O.; Project Administration, J.M.O.

## Additional Information

The authors declare no competing interests.

