## Supplementary Information - Figures for "Multi-channel recordings reveal age-related differences in the sleep of juvenile and adult zebra finches"

### Supplementary Figures

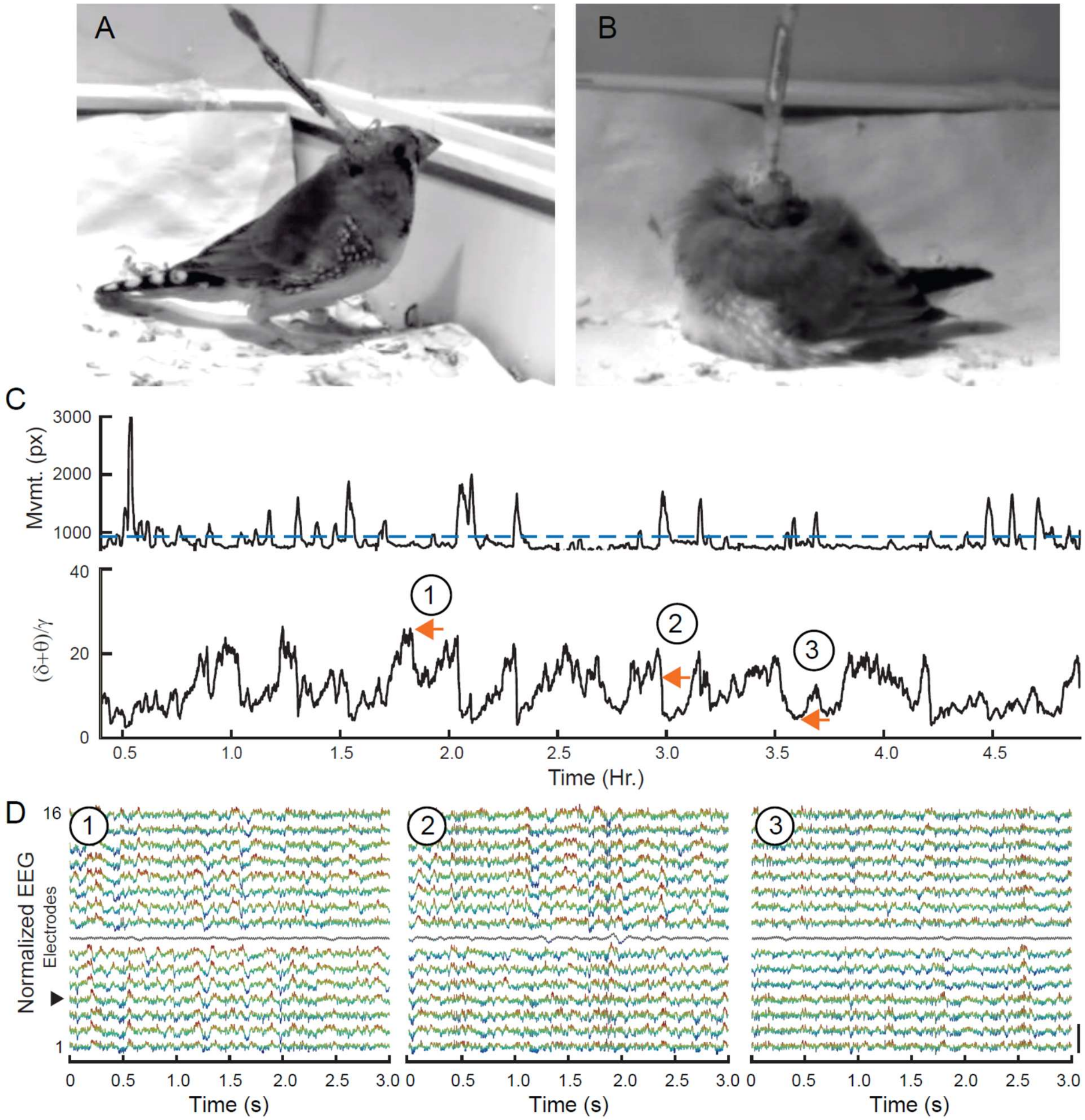


**Figure S1. Multichannel EEG recordings during sleep in a juvenile.** (A) Image of an awake tethered juvenile bird. (B) Image of a sleeping tethered juvenile. Same bird as in (A). Note how bird is able to assume a sleep-specific posture despite the tether cable. (C) Top: Bird movement extracted from the infrared video recording (px, pixels). Blue dashed line indicates the threshold delineating wake and sleep. Bottom: Corresponding $(\delta+\theta)/\gamma$ trace shows the oscillatory components of EEG, representing different sleep stages. Orange arrows 1, 2, and 3, correspond to EEG data in (D). Movement and $(\delta+\theta)/\gamma$ traces are smoothed with a 30s window for visualization purposes. (D) 3s examples of simultaneous EEG recorded from the 16 different electrodes (bottom trace, electrode 1; top trace, electrode 16). Circled numbers correspond to orange arrows in (B) and indicate examples of SWS (1), IS (2), and REM sleep (3). Color scheme is for visualization purposes only. Black arrow indicates electrode 4. Electrode 8 (grey line) was noisy and discarded in further analysis.

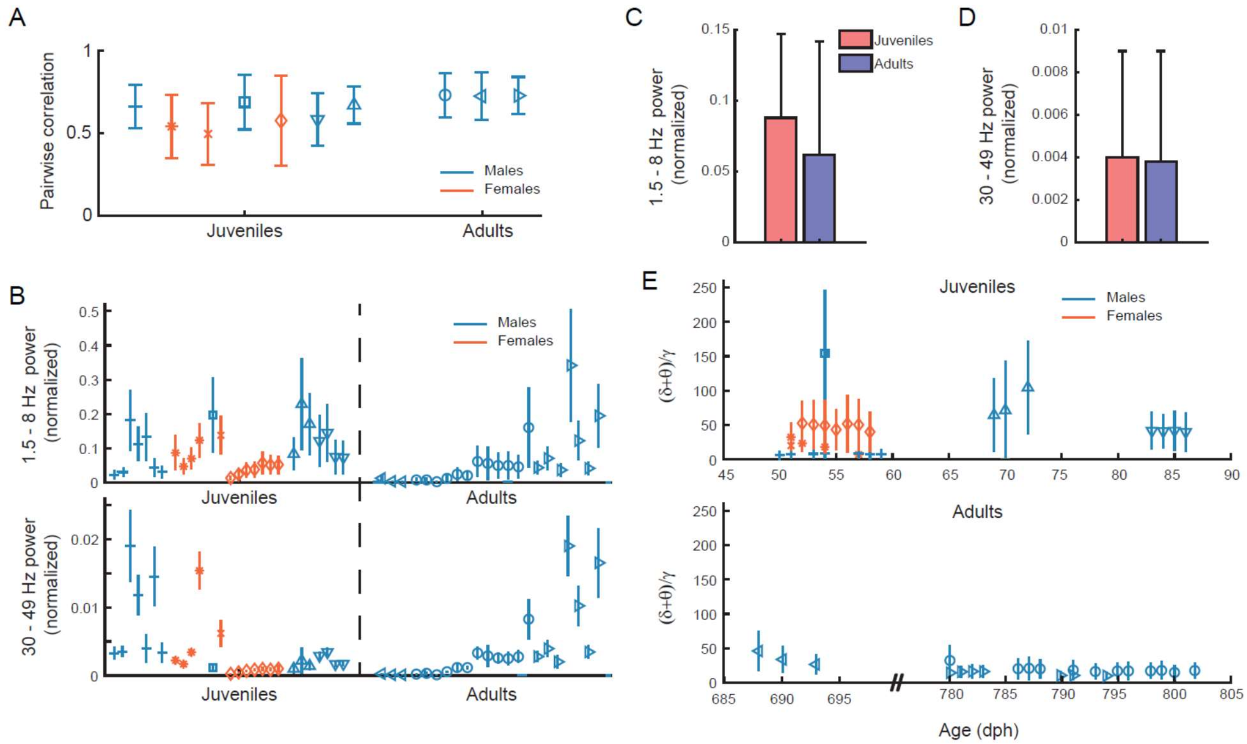


**Figure S2. Low and high frequency components of the $(\delta+\theta)/\gamma$ trace in adults and juveniles.** (A) Mean correlation calculated for $(\delta+\theta)/\gamma$ traces across all electrode pairs for the first recording night. Each symbol represents a different bird; blue symbols indicate males; orange symbols indicate females. Error bars indicate SD. (B) $(\delta+\theta)/\gamma$ values were significantly larger in juveniles compared to adults. We examined the power in each of these bands separately for juveniles and adults. Top: $(\delta+\theta)$ values calculated across 12 hours of sleep in 3 s bins, for all nights of each bird (median ± interquartile). Each symbol represents a different bird, and different nights of sleep are presented sequentially. Blue symbols indicate males; orange symbols indicate females. Bottom: $\gamma$ values calculated across 12 hours of sleep in 3s bins, for all nights of each bird. (C) Mean relative power for the $(\delta+\theta)$ band was larger in juveniles compared to adults, but this difference was not significant (p=0.07; two-way unbalanced ANOVA). (D). Mean relative power for the $\gamma$ band was not significantly different between juveniles and adults (p=0.47; two-way unbalanced ANOVA). (E) $(\delta+\theta)/\gamma$ values are plotted as function of age for juvenile birds (upper plot) and adults (lower plot). Data are the same as plotted in Fig.1E. Figure conventions same as in (B).

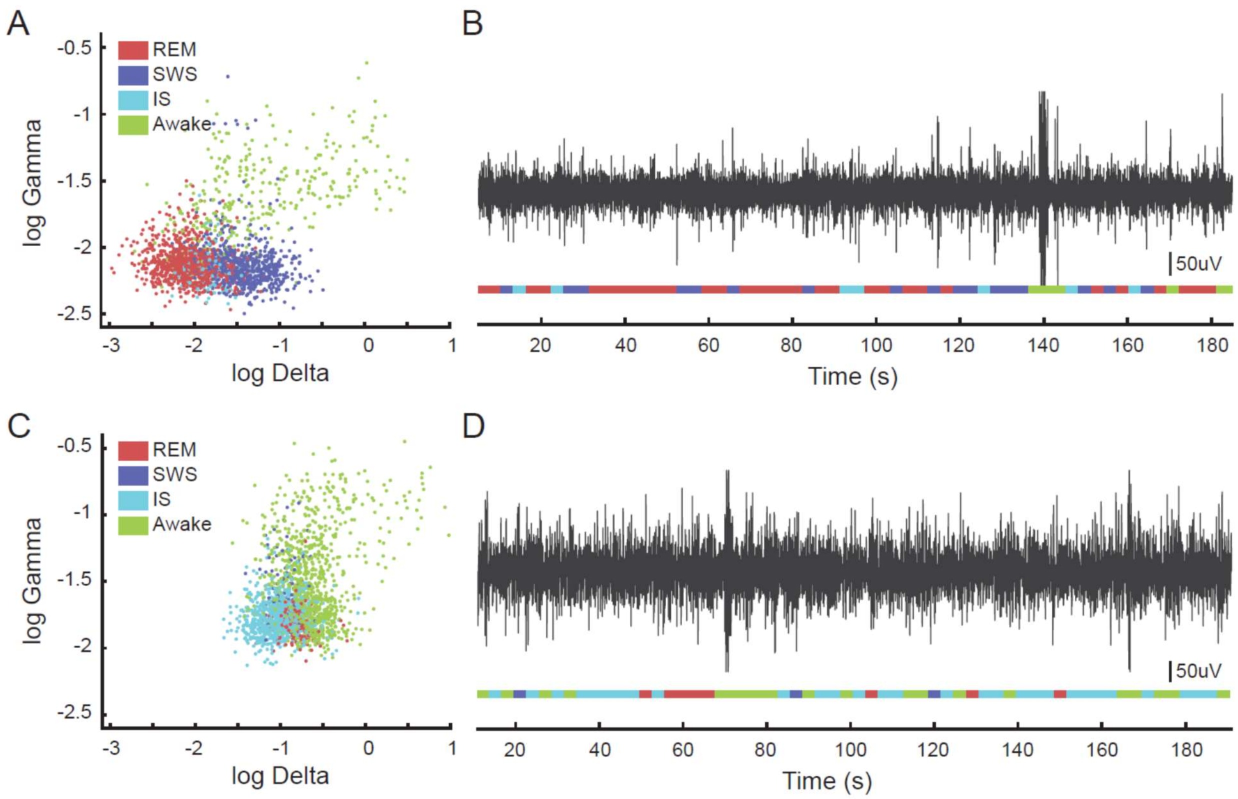


**Figure S3. Examples of successful and unsuccessful automatic segmentation of sleep stages.** (A) Clustering output for a successful segmentation (from adult 72-00). Note how clusters are separated in the log gamma and log delta space. (B) 3-minute example trace of EEG that is labeled according to clustering in (A). (C) Clustering output for an unsuccessful segmentation (from juvenile w018; none of the nights could be clustered for this bird). Note how clusters overlap heavily in the log gamma and log delta space. (D). 3-minute example trace of EEG that is labeled according to clustering in (C). Note how numerous segments are labeled as Awake or IS.

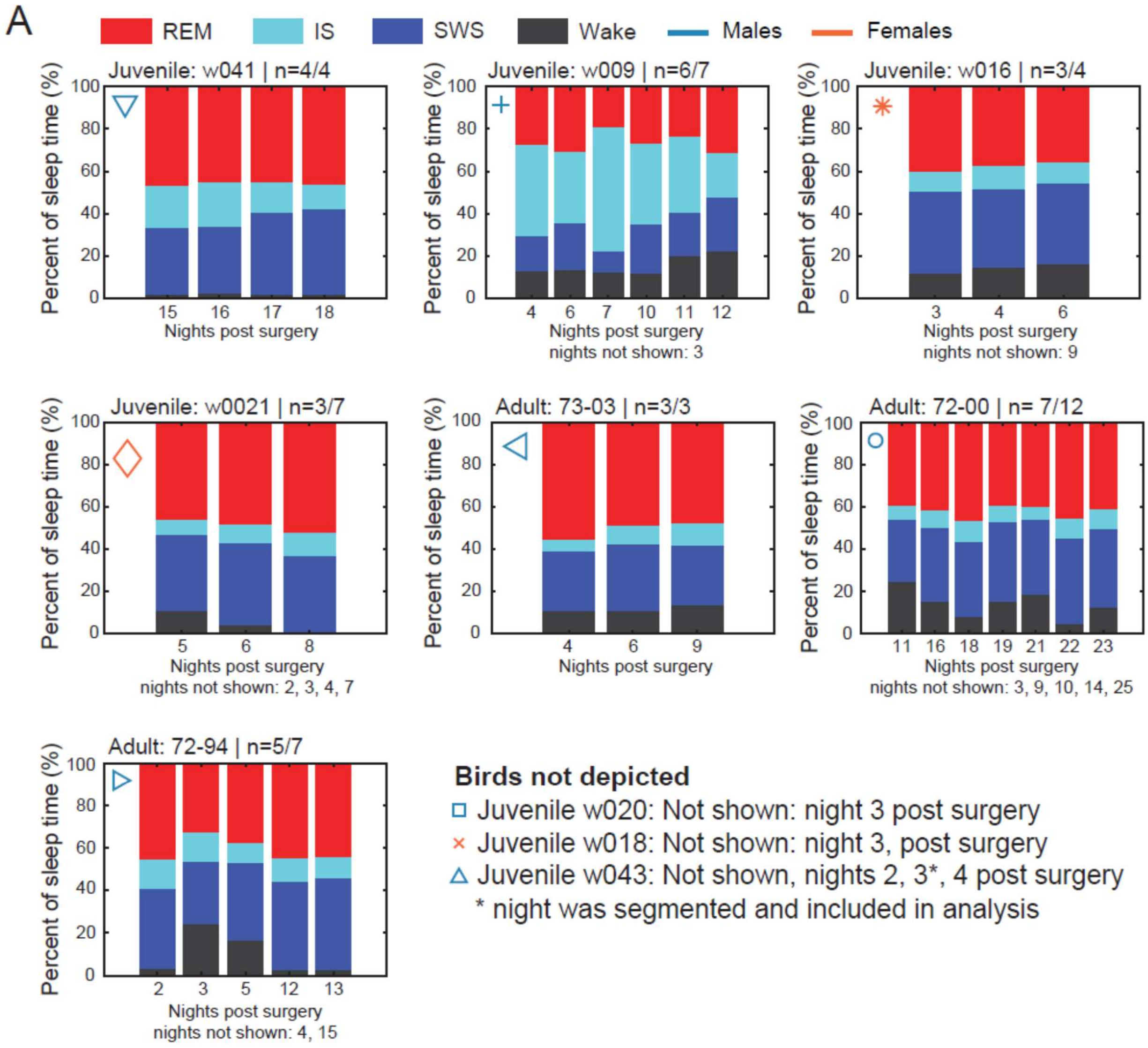


**Figure S4. Percentage of sleep stages across nights for each bird.** (A) Stacked plots represent the percentage of each sleep stage across recording nights. Each panel contains the data from an individual bird; the symbol in the upper left corner matches the identification of the bird in Fig. 1E. The sleep scoring used in this plot is the result of an automatic clustering algorithm.[1,2] Only one night of sleep was segmented for juvenile bird w043 (Δ) and therefore it is not included here. The number of successfully clustered nights out of the total number of nights are indicated in the title for each panel (e.g., n=4/4), and the specific nights that were not successfully clustered are indicated as "nights not shown". Note how the percentage of sleep across nights for individual birds is largely consistent across nights.

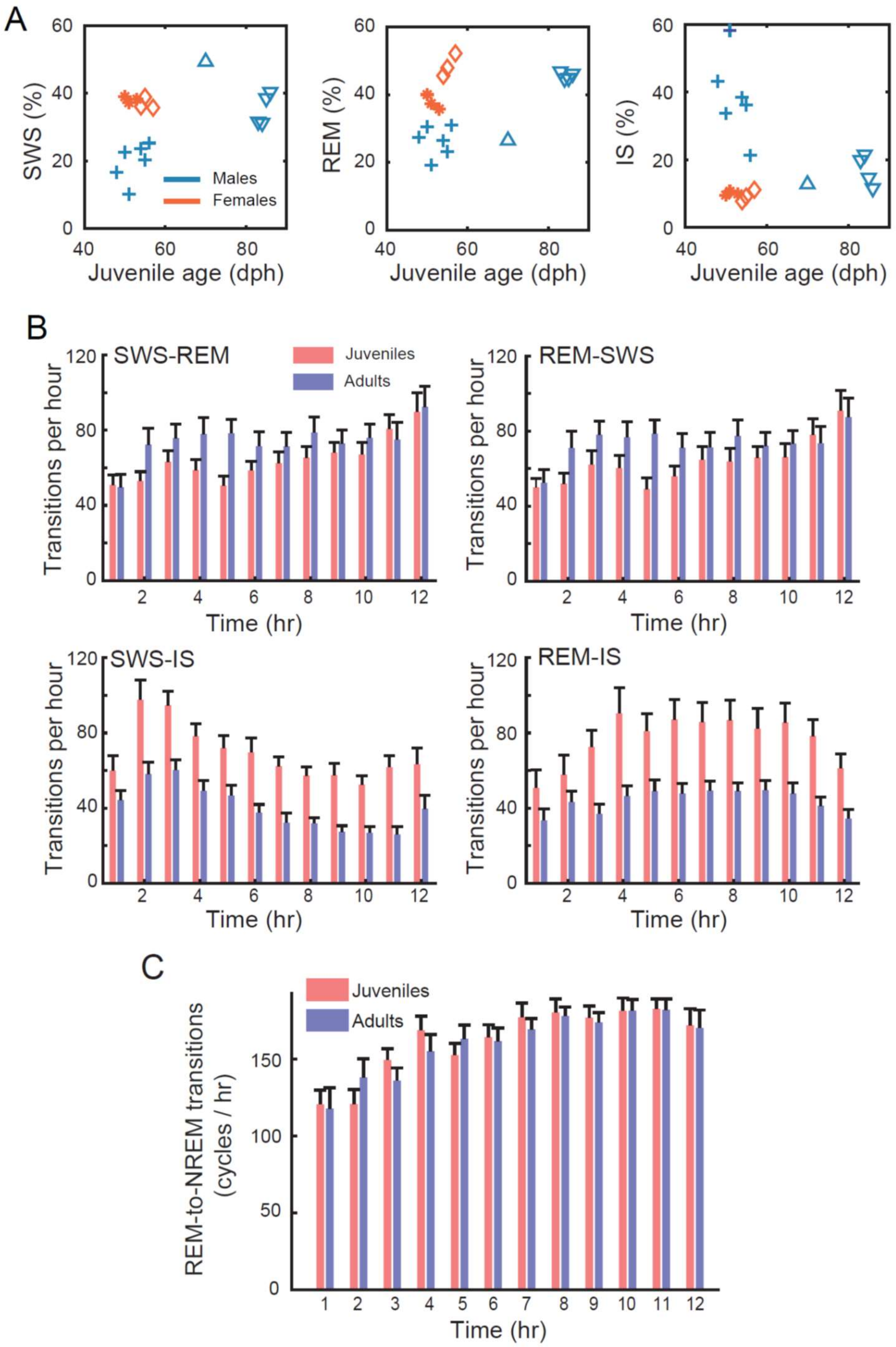


**Figure S5 Analysis of REM versus NREM sleep in adults and juveniles** (A)Scatter plots depict the percentage of SWS, REM sleep, or IS per night as a function of juvenile age. Same animals as in Fig. 2E. A clear linear relationship between sleep stage percentage and age is not present. (B) Bar plots indicate the average number of transitions per hour from SWS to REM, REM to SWS, SWS to IS, and REM to IS over 12 hours of sleep for juveniles (red) and adults (blue). Error bars indicated the s.e.m. (C) REM to NREM (IS+SWS) for juveniles (red) and adults (blue) over 12 hours of sleep. Error bars indicated the s.e.m.

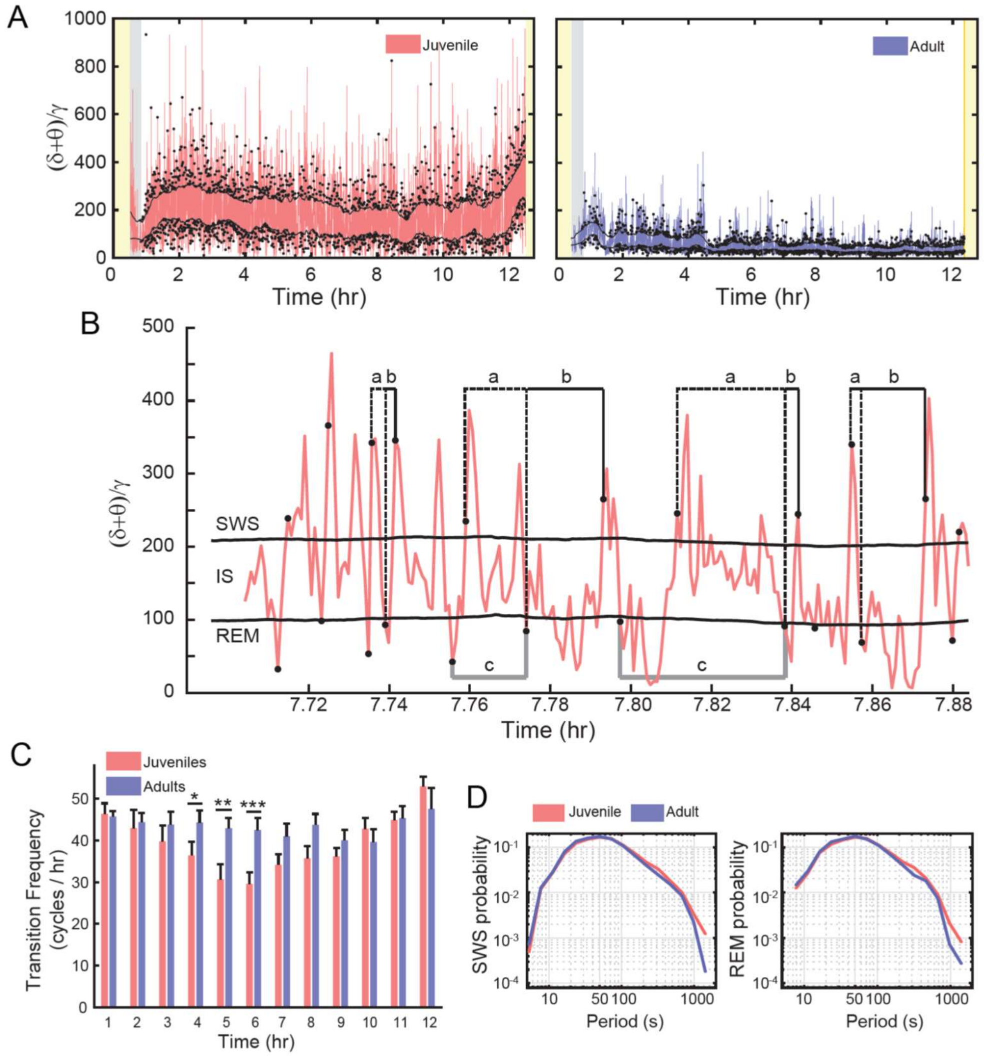


**Figure S6 State transitions during continuous sleep (**A) Unsmoothed $(\delta+\theta)/\gamma$ traces for a juvenile bird (left) and an adult bird (right) for 12 hours of sleep (same birds as depicted in Fig. 1D and indicated as black arrows in Fig. 1E). In each $(\delta+\theta)/\gamma$ trace, the upper and lower black lines indicate the SWS and REM detection threshold, respectively. Gray dots indicate the first detection after a threshold crossing (top dots, SWS; bottom dots, REM). The yellow bars indicated periods when the light was on. Gray shading indicates time periods after the lights were off that the birds were still awake based on movement detection. (B) Enlargement of a short window of the unsmoothed $(\delta+\theta)/\gamma$ trace from the juvenile in (A). Dotted black lines labeled ‘a’ indicate examples of detected SWS to REM transitions. Note how the duration of ‘a’ varies as a function of how much time is spent in SWS and IS. Solid black lines labeled ‘b’ indicate examples of REM to SWS transitions; the combined duration of ‘a’ plus ‘b’ completes one full cycle of SWS (SWS-REM-SWS). Solid gray lines labeled ‘c’ indicate examples of one full cycle of REM (REM-SWS-REM). (C) Bar plot indicates the average number of SWS-to-REM transitions (i.e., the number of transitions labeled as ‘a’ in (B) per hour during night in juveniles and adults. For adults, the transitions were relatively constant, whereas for juveniles, the transition frequency varied throughout the night (see text for details). The transition frequency was significantly lower for juveniles compared to adults for the 5$^{th}$, 6$^{th}$, and 7$^{th}$ hours. (D) The probability distribution functions for SWS cycles (left plot; SWS cycle indicated as ‘a’ plus ‘b’ in (B)) and REM cycles (right plot; REM cycle indicated as ‘c’ in (B)). The probability of observing long SWS and REM cycles is significantly higher in the juvenile group. SWS cycle was 92.86 ± 121.41 s in juveniles and 82.84 ± 102.80 s in adults (unpaired t-test, $p=5.7\times10^{-12}$). Similarly, REM cycles were also significantly longer in the juveniles (92.88 ± 115.07 s in juveniles, and 82.74 ± 99.15 s in adults; unpaired t-test, $p=5.5\times10^{-13}$).

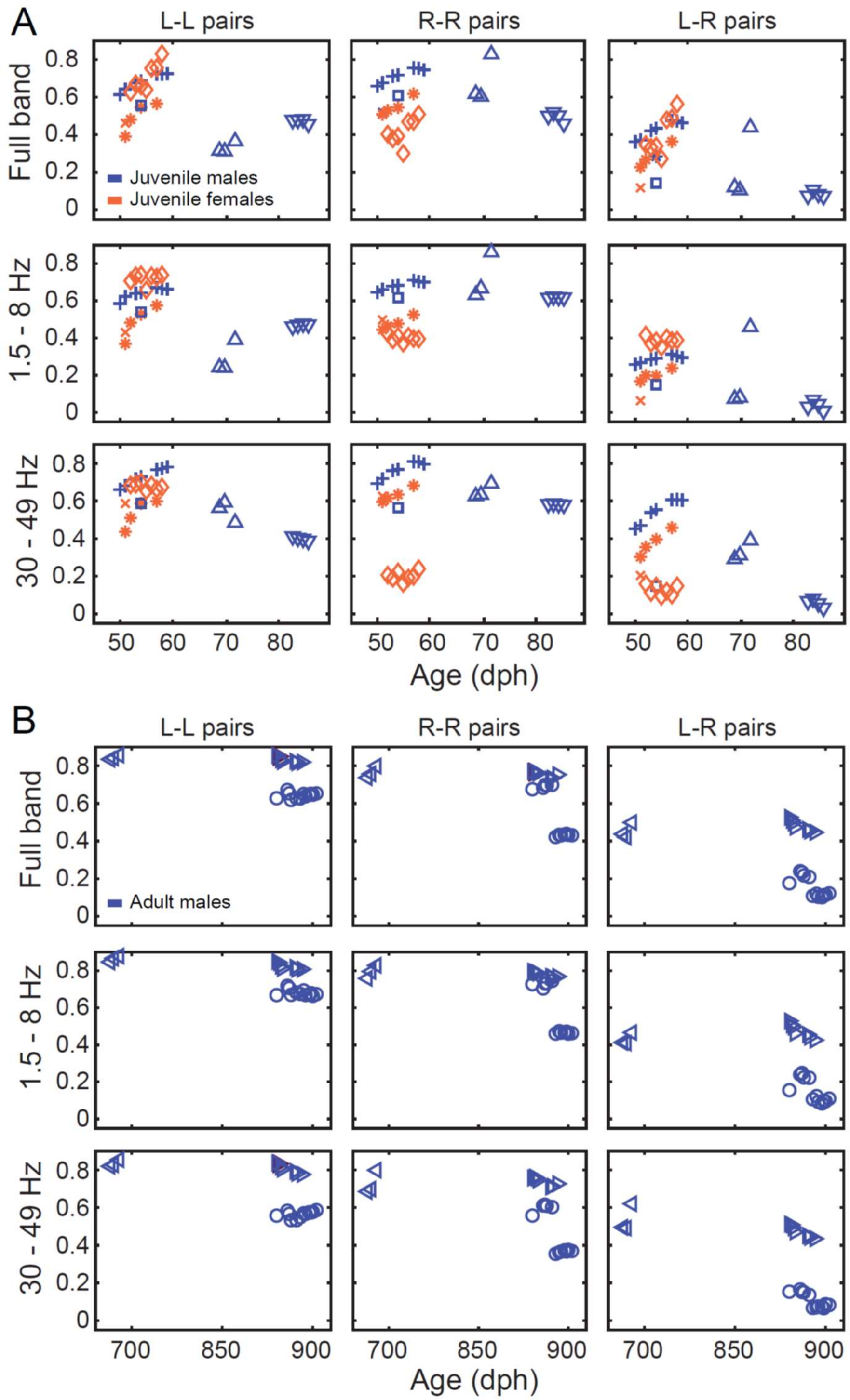


**Figure S7**. **Left intra-hemispheric functional connectivity increases linearly during early maturation in juveniles, but stabilizes or declines by approaching maturity.** (A) The increasing linear relationship between connectivity and age was only observed in younger juveniles (the 5 birds presented in Fig. 4B). Panels display the data from all 7 juveniles. Same figure convention as for Fig. 4 Note how an increasing linear trend does not exist for older juveniles. (B). This data supports the view that the connectivity elevates at a high pace in early juvenile stage and then stabilizes or declines before the age of maturation to the adulthood level.

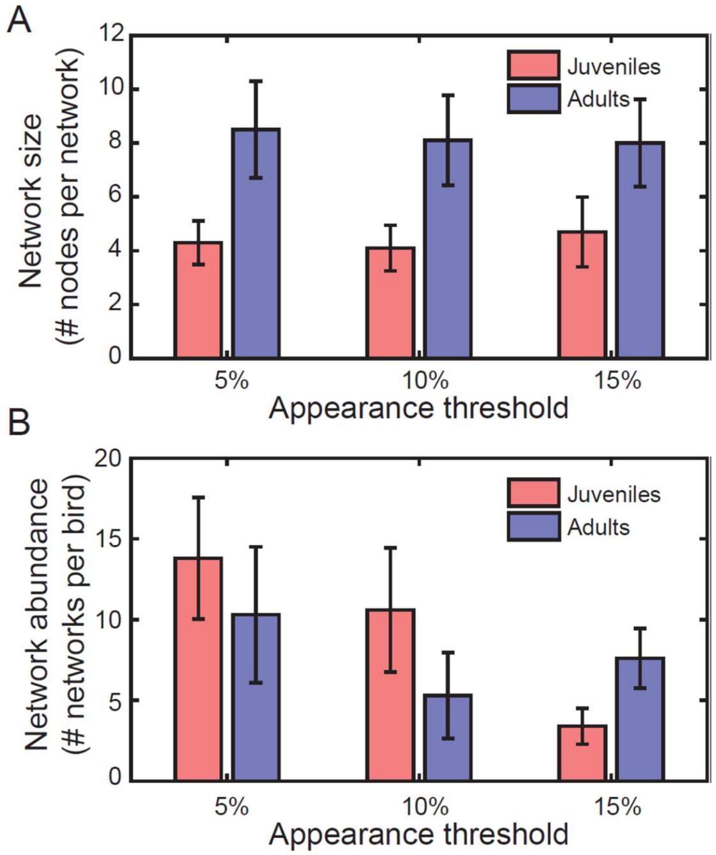


**Figure S8 Effects of setting different appearance thresholds on the network analysis results.** (A) The appearance threshold (x axis) is a parameter that we used to constrain the network analysis, such that a dominant network had to appear for at least 10% of the bins throughout the night, for three separate nights.
