## Supplementary Information - ANOVA Statistics for "Multi-channel recordings reveal age-related differences in the sleep of juvenile and adult zebra finches"

### Analysis of Variance tables for the tests presented in the main text

Two-way ANOVA for **$(\delta+\theta)/\gamma$**, x1: age, x2: sex, using anovan MATLAB function

| Source | Sum Sq. | d.f. | Mean Sq. | F | Prob>F |
|---|---|---|---|---|---|
| X1 | 4138.6488 | 1 | 4138.6488 | 6.2945 | 0.032698 |
| X2 | 141.6964 | 1 | 141.6964 | 0.21551 | 0.64568 |
| Error | 30245.2729 | 46 | 657.5059 | | |
| Total | 34977.0134 | 48 | | | |

Constrained (Type III) sums of squares.

Two-way ANOVA for **$(\delta+\theta)$**, x1: age, x2: sex, using anovan MATLAB function

| Source | Sum Sq. | d.f. | Mean Sq. | F | Prob>F |
|---|---|---|---|---|---|
| X1 | 0.015669 | 1 | 0.015669 | 3.2951 | 0.076009 |
| X2 | 0.0084312 | 1 | 0.0084312 | 1.773 | 0.18957 |
| Error | 0.21874 | 46 | 0.0047553 | | |
| Total | 0.23547 | 48 | | | |

Constrained (Type III) sums of squares.

Two-way ANOVA for $\gamma$, x1: age, x2: sex, using anovan MATLAB function

| Source | Sum Sq. | d.f. | Mean Sq. | F | Prob>F |
|---|---|---|---|---|---|
| X1 | 1.3207e-05 | 1 | 1.3207e-05 | 0.51222 | 0.4778 |
| X2 | 3.5236e-05 | 1 | 3.5236e-05 | 1.3665 | 0.24843 |
| Error | 0.0011861 | 46 | 2.5784e-05 | | |
| Total | 0.0012218 | 48 | | | |

Constrained (Type III) sums of squares.

Two-way ANOVA for **REM percentage**, x1: age, x2: sex, using anovan MATLAB function

| Source | Sum Sq. | d.f. | Mean Sq. | F | Prob>F |
|---|---|---|---|---|---|
| X1 | 620.137 | 1 | 620.137 | 10.7135 | 0.0027508 |
| X2 | 372.7795 | 1 | 372.7795 | 6.4402 | 0.01679 |
| Error | 1678.6209 | 29 | 57.8835 | | |
| Total | 2380.5423 | 31 | | | |

Constrained (Type III) sums of squares.

Two-way ANOVA for **SWS percentage**, x1: age, x2: sex, using anovan MATLAB function

| Source | Sum Sq. | d.f. | Mean Sq. | F | Prob>F |
|---|---|---|---|---|---|
| X1 | 308.5916 | 1 | 308.5916 | 5.4826 | 0.026288 |
| X2 | 342.2265 | 1 | 342.2265 | 6.0801 | 0.019828 |
| Error | 1632.2957 | 29 | 56.2861 | | |
| Total | 2081.2443 | 31 | | | |

Constrained (Type III) sums of squares.

Two-way ANOVA for **IS percentage**, x1: age, x2: sex, using anovan MATLAB function

| Source | Sum Sq. | d.f. | Mean Sq. | F | Prob>F |
|---|---|---|---|---|---|
| X1 | 2243.3562 | 1 | 2243.3562 | 28.6299 | 9.573e-06 |
| X2 | 1344.5007 | 1 | 1344.5007 | 17.1586 | 0.0002715 |
| Error | 2272.3579 | 29 | 78.3572 | | |
| Total | 4809.4547 | 31 | | | |

Constrained (Type III) sums of squares.

Two-way ANOVA for **REM duration**, x1: age, x2: sex, using anovan MATLAB function

| Source | Sum Sq. | d.f. | Mean Sq. | F | Prob>F |
|---|---|---|---|---|---|
| X1 | 15.4459 | 1 | 15.4459 | 6.3213 | 0.017732 |
| X2 | 13.1744 | 1 | 13.1744 | 5.3917 | 0.027459 |
| Error | 70.8609 | 29 | 2.4435 | | |
| Total | 90.6333 | 31 | | | |

Constrained (Type III) sums of squares.

Two-way ANOVA for **SWS duration**, x1: age, x2: sex, using anovan MATLAB function

| SourceSum | Sq. | d.f. | Mean Sq. | F | Prob>F |
|---|---|---|---|---|---|
| X1 | 14.0336 | 1 | 14.0336 | 11.7447 | 0.0018455 |
| X2 | 14.2408 | 1 | 14.2408 | 11.918 | 0.0017279 |
| Error | 34.6518 | 29 | 1.1949 | | |
| Total | 54.1352 | 31 | | | |

Constrained (Type III) sums of squares.

Two-way ANOVA for **IS duration**, x1: age, x2: sex, using anovan MATLAB function

| SourceSum | Sq. | d.f. | Mean Sq. | F | Prob>F |
|---|---|---|---|---|---|
| X1 | 14.4968 | 1 | 14.4968 | 6.7135 | 0.014823 |
| X2 | 11.7487 | 1 | 11.7487 | 5.4409 | 0.026817 |
| Error | 62.6208 | 29 | 2.1593 | | |
| Total | 80.7874 | 31 | | | |

Constrained (Type III) sums of squares.

One-way ANOVA for **REM Percentage**, x1: juvenile sex, using anovan MATLAB function

| Source | Sum Sq. | d.f. | Mean Sq. | F | Prob>F |
|---|---|---|---|---|---|
| X1 | 372.78 | 1 | 372.78 | 4.42 | 0.0529 |
| Error | 1265.7 | 15 | 84.38 | | |
| Total | 1638.48 | 16 | | | |

One-way ANOVA for **SWS Percentage**, x1: juvenile sex, using anovan MATLAB function

| Source | Sum Sq. | d.f. | Mean Sq. | F | Prob>F |
|---|---|---|---|---|---|
| X1 | 342.23 | 1 | 342.227 | 3.91 | 0.0667 |
| Error | 1312.77 | 15 | 87.518 | | |
| Total | 1655 | 16 | | | |

One-way ANOVA for **IS Percentage**, x1: juvenile sex, using anovan MATLAB function

| Source | Sum Sq. | d.f. | Mean Sq. | F | Prob>F |
|---|---|---|---|---|---|
| X1 | 1344.5 | 1 | 1344.5 | 9.21 | 0.0084 |
| Error | 2189.49 | 15 | 145.97 | | |
| Total | 3533.99 | 16 | | | |

One-way ANOVA for **REM Duration**, x1: juvenile sex, using anovan MATLAB function

| Source | Sum Sq. | d.f. | Mean Sq. | F | Prob>F |
|---|---|---|---|---|---|
| X1 | 21.3391 | 1 | 21.3391 | 54.31 | 2.33001e-06 |
| Error | 5.8933 | 15 | 0.3929 | | |
| Total | 27.2324 | 16 | | | |

One-way ANOVA for **SWS Duration**, x1: juvenile sex, using anovan MATLAB function

| Source | Sum Sq. | d.f. | Mean Sq. | F | Prob>F |
|---|---|---|---|---|---|
| X1 | 28.5115 | 1 | 28.5115 | 117.58 | 1.70414e-08 |
| Error | 3.6371 | 15 | 0.2425 | | |
| Total | 32.1486 | 16 | | | |

One-way ANOVA for **IS Duration**, x1: juvenile sex, using anovan MATLAB function

| Source | Sum Sq. | d.f. | Mean Sq. | F | Prob>F |
|---|---|---|---|---|---|
| X1 | 12.4023 | 1 | 12.4023 | 34.76 | 2.93802e-05 |
| Error | 5.3517 | 15 | 0.3568 | | |
| Total | 17.754 | 16 | | | |

Two-way ANOVA for **local wave occurrence rate**, x1: age, x2: sex, using anovan MATLAB function

| Source | Sum Sq. | d.f. | Mean Sq. | F | Prob>F |
|---|---|---|---|---|---|
| X1 | 0.78796 | 1 | 0.78796 | 25.1373 | 6.7767e-07 |
| X2 | 0.19424 | 1 | 0.19424 | 6.1967 | 0.013031 |
| Error | 21.8798 | 698 | 0.031346 | | |
| Total | 23.6767 | 700 | | | |

Constrained (Type III) sums of squares.

Two-way ANOVA for **connectivity across sleep stages in adults**, x1: [SWS, IS, REM stages], x2: [LL, RR, LR connectivity], using anova2 MATLAB function

| Source | SS | df | MS | F | Prob>F |
|---|---|---|---|---|---|
| Columns | 7.0207 | 2 | 3.5103 | 239.7073 | 1.4414e-52 |
| Rows | 0.075048 | 2 | 0.037524 | 2.5624 | 0.079799 |
| Interaction | 0.054864 | 4 | 0.013716 | 0.93661 | 0.44387 |
| Error | 2.7678 | 189 | 0.014644 | | |
| Total | 9.9184 | 197 | | | |

Two-way ANOVA for **connectivity across sleep stages in juveniles**, x1: [SWS, IS, REM stages], x2: [LL, RR, LR connectivity], using anova2 MATLAB function

| Source | SS | df | MS | F | Prob>F |
|---|---|---|---|---|---|
| Columns | 0.021983 | 2 | 0.010991 | 0.53371 | 0.5874 |
| Rows | 2.8498 | 2 | 1.4249 | 69.1902 | 9.6389e-23 |
| Interaction | 0.0043447 | 4 | 0.0010862 | 0.052742 | 0.99476 |
| Error | 3.5216 | 171 | 0.020594 | | |
| Total | 6.3978 | 179 | | | |

Two-way ANOVA for **REM-to-IS** transitions, x1: age, x2: sex, using anovan MATLAB function

| Source | Sum Sq. | d.f. | Mean Sq. | F | Prob>F |
|---|---|---|---|---|---|
| X1 | 2326266.7508 | 1 | 2326266.7508 | 27.0961 | 1.3043e-05 |
| X2 | 1315662.4251 | 1 | 1315662.4251 | 15.3247 | 0.00048249 |
| Error | 2575573.0455 | 30 | 85852.4348 | | |
| Total | 5157324.2424 | 32 | | | |

Two-way ANOVA for **REM-to-SWS** transitions, x1: age, x2: sex, using anovan MATLAB function

| Source | Sum Sq. | d.f. | Mean Sq. | F | Prob>F |
|---|---|---|---|---|---|
| X1 | 390367.1928 | 1 | 390367.1928 | 4.8759 | 0.03502 |
| X2 | 454779.787 | 1 | 454779.787 | 5.6804 | 0.023687 |
| Error | 2401833.4924 | 30 | 80061.1164 | | |
| Total | 2983183.5152 | 32 | | | |

Two-way ANOVA for **SWS-to-IS** transitions, x1: age, x2: sex, using anovan MATLAB function

| Source | Sum Sq. | d.f. | Mean Sq. | F | Prob>F |
|---|---|---|---|---|---|
| X1 | 1556397.3064 | 1 | 1556397.3064 | 46.906 | 1.3373e-07 |
| X2 | 623673.0918 | 1 | 623673.0918 | 18.796 | 0.0001508 |
| Error | 995435.3788 | 30 | 33181.1793 | | |
| Total | 2612580.2424 | 32 | | | |

Two-way ANOVA for **SWS-to-REM** transitions, x1: age, x2: sex, using anovan MATLAB function

| Source | Sum Sq. | d.f. | Mean Sq. | F | Prob>F |
|---|---|---|---|---|---|
| X1 | 341905.4848 | 1 | 341905.4848 | 4.8873 | 0.034822 |
| X2 | 347332.5143 | 1 | 347332.5143 | 4.9649 | 0.033512 |
| Error | 2098746.0152 | 30 | 69958.2005 | | |
| Total | 2571704.2424 | 32 | | | |

One-way ANOVA for **SWS-to-REM** transitions, x1: juvenile sex, using anovan MATLAB function

| Source | Sum Sq. | d.f. | Mean Sq. | F | Prob>F |
|---|---|---|---|---|---|
| X1 | 2412.03 | 1 | 2412.03 | 8.41 | 0.011 |
| Error | 4299.78 | 15 | 286.65 | | |
| Total | 6711.82 | 16 | | | |

One-way ANOVA for **REM-to-SWS** transitions, x1: juvenile sex, using anovan MATLAB function

| Source | Sum Sq. | d.f. | Mean Sq. | F | Prob>F |
|---|---|---|---|---|---|
| X1 | 3158.19 | 1 | 3158.19 | 7.42 | 0.0157 |
| Error | 6380.4 | 15 | 425.36 | | |
| Total | 9538.59 | 16 | | | |

One-way ANOVA for **REM-to-IS** transitions, x1: juvenile sex, using anovan MATLAB function

| Source | Sum Sq. | d.f. | Mean Sq. | F | Prob>F |
|---|---|---|---|---|---|
| X1 | 9136.5 | 1 | 9136.54 | 9.82 | 0.0068 |
| Error | 13956.2 | 15 | 930.41 | | |
| Total | 23092.7 | 16 | | | |

One-way ANOVA for **SWS-to-IS** transitions, x1: juvenile sex, using anovan MATLAB function

| Source | Sum Sq. | d.f. | Mean Sq. | F | Prob>F |
|---|---|---|---|---|---|
| X1 | 4331.06 | 1 | 4331.06 | 20.14 | 0.0004 |
| Error | 3226.43 | 15 | 215.1 | | |
| Total | 7557.49 | 16 | | | |

Two-way ANOVA for **CC** in **LL, RR,** and **LR** pair of channels, in **adults**, x1: SWS, IS, REM, x2: LL, RR, and LR, using anova2 MATLAB function

| Source | SS | df | MS | F | Prob>F |
|---|---|---|---|---|---|
| Columns | 7.0207 | 2 | 3.5103 | 239.7073 | 1.4414e-52 |
| Rows | 0.075048 | 2 | 0.037524 | 2.5624 | 0.079799 |
| Interaction | 0.054864 | 4 | 0.013716 | 0.93661 | 0.44387 |
| Error | 2.7678 | 189 | 0.014644 | | |
| Total | 9.9184 | 197 | | | |

Accompanying pairwise comparisons from Tukey multiple comparison test (multcompare function):

| Group1 | Group2 | p-value |
|---|---|---|
| LL | RR | 0.0396 |

| | | |
|---|---|---|
| LL | LR | 9.56e-10 |
| RR | LR | 9.56e-10 |

Two-way ANOVA for **CC** in **LL, RR,** and **LR** pair of channels, in **juveniles**, x1: SWS, IS, REM, x2: LL, RR, and LR, using anova2 MATLAB function

| Source | SS | df | MS | F | Prob>F |
|---|---|---|---|---|---|
| Columns | 2.8498 | 2 | 1.4249 | 69.1902 | 9.6389e-23 |
| Rows | 0.021983 | 2 | 0.010991 | 0.53371 | 0.5874 |
| Interaction | 0.0043447 | 4 | 0.0010862 | 0.052742 | 0.99476 |
| Error | 3.5216 | 171 | 0.020594 | | |
| Total | 6.3978 | 179 | | | |

Accompanying pairwise comparisons from Tukey multiple comparison test (multcompare function):

| Group1 | Group2 | p-value |
|---|---|---|
| LL | RR | 0.5489 |
| LL | LR | 9.56e-10 |
| RR | LR | 9.56e-10 |
